# Effect of obesity on the acute response to SARS-CoV-2 infection and development of post-acute sequelae of COVID-19 (PASC) in nonhuman primates

**DOI:** 10.1101/2025.02.18.638792

**Authors:** Kristin A. Sauter, Gabriela M. Webb, Lindsay Bader, Craig N. Kreklywich, Diana L. Takahashi, Cicely Zaro, Casey M. McGuire, Anne D. Lewis, Lois M. A. Colgin, Melissa A. Kirigiti, Hannah Blomenkamp, Cleiton Pessoa, Matthew Humkey, Jesse Hulahan, Madeleine Sleeman, Robert C. Zweig, Sarah Thomas, Archana Thomas, Lina Gao, Alec. J. Hirsch, Mayaan Levy, Sara R. Cherry, Steven E. Kahn, Mark K. Slifka, Daniel N. Streblow, Jonah B. Sacha, Paul Kievit, Charles T. Roberts

## Abstract

Long-term adverse consequences of SARS-CoV-2 infection, termed “long COVID” or post-acute sequelae of COVID (PASC), are a major component of overall COVID-19 disease burden. Prior obesity and metabolic disease increase the severity of acute disease, but SARS-CoV-2 infection also contributes to the development of new-onset metabolic disease. Since the COVID pandemic occurred in the context of the global obesity epidemic, an important question is the extent to which pre-existing obesity modifies long-term responses to SARS-CoV-2 infection. We utilized a nonhuman primate model to compare the effects of infection with the SARS-CoV-2 delta variant in lean and obese/insulin-resistant adult male rhesus macaques over a 6-month time course. While some longitudinal responses to SARS-CoV-2 infection, including overall viral dynamics, SARS-CoV-2-specific IgG induction, cytokine profiles, and tissue persistence of viral RNA, did not appreciably differ between lean and obese animals, other responses, including neutralizing Ab dynamics, lung pathology, body weight, degree of insulin sensitivity, adipocytokine profiles, body temperature, and nighttime activity levels were significantly different in lean versus obese animals. Furthermore, several parameters in lean animals were altered following SARS-CoV-2 infection to resemble those in obese animals. Notably, persistent changes in multiple parameters were present in most animals, suggesting that PASC may be more prevalent than estimated from self-reported symptoms in human studies.

## Introduction

The COVID-19 pandemic has caused significant global morbidity and mortality and remains a persistent public health issue due to the emergence of variants of concern and the expanding spectrum of post-acute sequelae of COVID-19 (PASC), also termed “long COVID” (1–5). An important feature of the pandemic was its appearance during the global obesity pandemic (6). COVID-19 severity is increased by comorbidities linked to obesity and insulin resistance (IR), including diabetes, cardiovascular disease, dyslipidemia, and hypertension (7–12), and obesity per se is an independent contributor to COVID-19 severity (13–19). This association is consistent with the adverse effects of obesity on the response to respiratory disease in general (20–23) and on viral respiratory disease in particular, including COVID-19 (24). Conversely, COVID-19 survivors often exhibit metabolic aspects of PASC that include dyslipidemia (12, 25, 26), altered glucose metabolism (27, 28), new-onset diabetes, and diabetic ketoacidosis (29–34), demonstrating that the relationship between COVID-19 and metabolic disease is reciprocal (35–37). Thus, COVID-19 severity is influenced by metabolic status, while COVID-19 itself influences the subsequent development of metabolic disease as an important component of PASC.

Since a principal aspect of obesity is an increase in white adipose tissue (WAT) mass, the potential role of WAT in SARS-CoV-2 infection outcomes is an important area of investigation (38). The increased WAT inflammation seen in COVID-19 patients (39) and the relationship between increased levels of the WAT-derived adipokine leptin and lung dysfunction (40) suggests that WAT itself may play a role in the effects of obesity on COVID-19 severity. Potential alterations in the expression of the adipokines leptin and adiponectin in SARS-CoV-2 infection (41–46) and subsequent effects on cardiometabolic risk (47, 48) may also contribute to the reciprocal interaction between SARS-CoV-2 and metabolic disease. Acute SARS-CoV-2 infection of WAT adipocytes and macrophages (49–52) suggests an additional mechanism through which obesity could influence COVID-19 severity; i.e., an increased number of cells susceptible to infection and subsequent pathologies such as dyslipidemia and increased inflammatory cytokine release and worsening insulin sensitivity. The overall scope of potential mechanisms through which altered WAT function contributes to metabolic aspects of PASC remains to be determined.

Animal models have proven invaluable in the investigation of COVID-19 disease (53) and multiple studies have investigated SARS-CoV-2 infection in nonhuman primate (NHP) species, including rhesus and cynomolgus macaques (52, 54–60). Macaques typically experience mild to moderate disease, but exhibit rapid increases in viral replication that last for several days, viral shedding for ∼2 weeks, and mild to moderate pulmonary disease, with interstitial pneumonia and accumulation of inflammatory macrophages, monocytes, and neutrophils in the lungs, and pulmonary immune responses to the virus (61, 62). In this study, we employed the rhesus macaque model to ascertain the potential effects of preexisting obesity on the acute and long-term response of multiple parameters to SARS-CoV-2 infection. We specifically wished to assess whether preexisting obesity and insulin resistance resulted in greater post-infection (PI) metabolic dysfunction in obese animals compared to that seen in lean, metabolically healthy animals (63–65). In addition to systemic metabolic parameters, we also assessed viral dynamics and SARS-CoV-2-specific serology, lung pathology, peripheral immune cell profiles, body composition, cytokine profiles, and circadian patterns of core body temperature (BT), and physical activity. We utilized the delta variant of SARS-CoV-2 in this study, since we (57) and others (66, 67) have previously demonstrated that this variant causes a disease phenotype in NHPs that mimics human COVID-19 pathology. Our studies with the delta variant also provide a necessary foundation for similar studies utilizing more recent omicron-related variants in the macaque model to identify intrinsic variant-specific differences in pathology that are not confounded by the variations in prior exposure, vaccination status, etc., present in clinical studies.

## Results

### Lean and obese groups exhibited distinct baseline phenotypes

For this study, obese animals had been maintained on an obesogenic western-style diet (WSD) for at least 12 months prior to assignment to the study. As shown in Table 1, there were no significant baseline differences in average age, although there was a similar wide range in each group. As expected, the lean and obese groups exhibited significant baseline differences in body weight (BW), % body fat, fat mass, lean mass, bone mass, bone mineral density (BMD), fasting blood insulin (FBI), Homeostatic Model Assessment for Insulin Resistance (HOMA-IR), hemoglobin A1c (HbA1c), total cholesterol (CHOL), HDL, and LDL. Fasting blood glucose (FBG) was similar in both groups. While there was a clear difference between group averages for triglycerides (TRIG), this was not statistically significant due to animal-to-animal variability. These data demonstrate that the obese group exhibited reduced insulin sensitivity but with relative euglycemia, indicated by similar FBG levels that were maintained by compensatory hyperinsulinemia reflected in the increased FBI values for the obese group.

**Table 1.** Baseline characteristics of lean and obese experimental groups. BMD, bone mineral density; FBG, fasting blood glucose; FBI, fasting blood insulin; HOMA-IR, Homeostasis Model Assessment for Insulin Resistance (FBGxFBI)/405; HbA1C, hemoglobin A1C; CHOL, cholesterol; TRIG, triglyceride. Data are presented as means ± SEM. Significance determined by unpaired 2-tailed t test. *, p<0.05; **, p<0.01; ***, p<0.001; ****, p<0.0001. NS, not significant.

|  | Lean | Obese | p value |
| --- | --- | --- | --- |
| N | 10 | 9 |  |
| Age (years) | 12.84 ± 1.42 (6.7-19.9) | 12.01 ± 1.83 (6.8-21.0) | NS |
| Weight (kg) | 10.03 ± 0.34 (8.20-11.65) | 16.54 ± 1.09 (12.75-21.25) | **** |
| Body Fat (%) | 11.81 ± 0.90 (7.59-16.89) | 33.64 ± 2.61 (20.94-48.03) | **** |
| Fat Mass (g) | 1121 ± 105.6 (689.5-1719) | 5624 ± 739.7 (2608-9715) | **** |
| Lean Mass (g) | 8170 ± 202.5 (7026-8996) | 10429 ± 411.9 (8680-12517) | **** |
| Bone Mass (g) | 416.7 ± 10.0 (372.0-484.7) | 477.7 ± 17.6 (397.9-549.1) | ** |
| BMD (g/cm<sup>2</sup>) | 0.673 ± 0.011 (0.613-0.734) | 0.732 ± 0.018 (0.648-0.824) | * |
| FBG (mg/dL) | 69.90 ± 3.32 (56-90) | 62.00 ± 2.48 (45-70) | NS |
| FBI (mU/ml) | 18.90 ± 4.76 (5.0-53.5) | 85.86 ± 18.23 (10.7-186.2) | ** |
| HOMA-IR | 3.52 ± 0.98 (0.69-10.43) | 12.81 ± 2.43 (1.54-26.49) | ** |
| HbA1C (%) | 4.46 ± 0.07 (4.1-4.8) | 4.87 ± 0.06 (4.6-5.2) | *** |
| CHOL (mg/dL) | 120.1 ± 4.4 (94-136) | 195.9 ± 15.2 (140-270) | *** |
| TRIG (mg/dL) | 42.60 ± 5.95 (18-71) | 113.30 ± 46.64 (26-483) | NS |
| HDL (mg/dL) | 64.60 ± 2.17 (53-75) | 95.44 ± 10.47 (37-150) | ** |
| LDL (mg/dL) | 47.00 ± 3.27 (27-63) | 67.56 ± 5.77 (38-99) | ** |

### SARS-Cov-2 infection differentially affects viral dynamics in lean and obese animals

To ascertain the effects of preexisting obesity and reduced insulin sensitivity on the acute and long-term responses to SARS-Cov-2 delta infection (B.1.617.2 strain), we infected 10 lean and 10 obese, insulin-resistant rhesus macaques and followed multiple viral, immune, and metabolic parameters for an additional 6 months as depicted in Figure 1.

**Figure 1.**
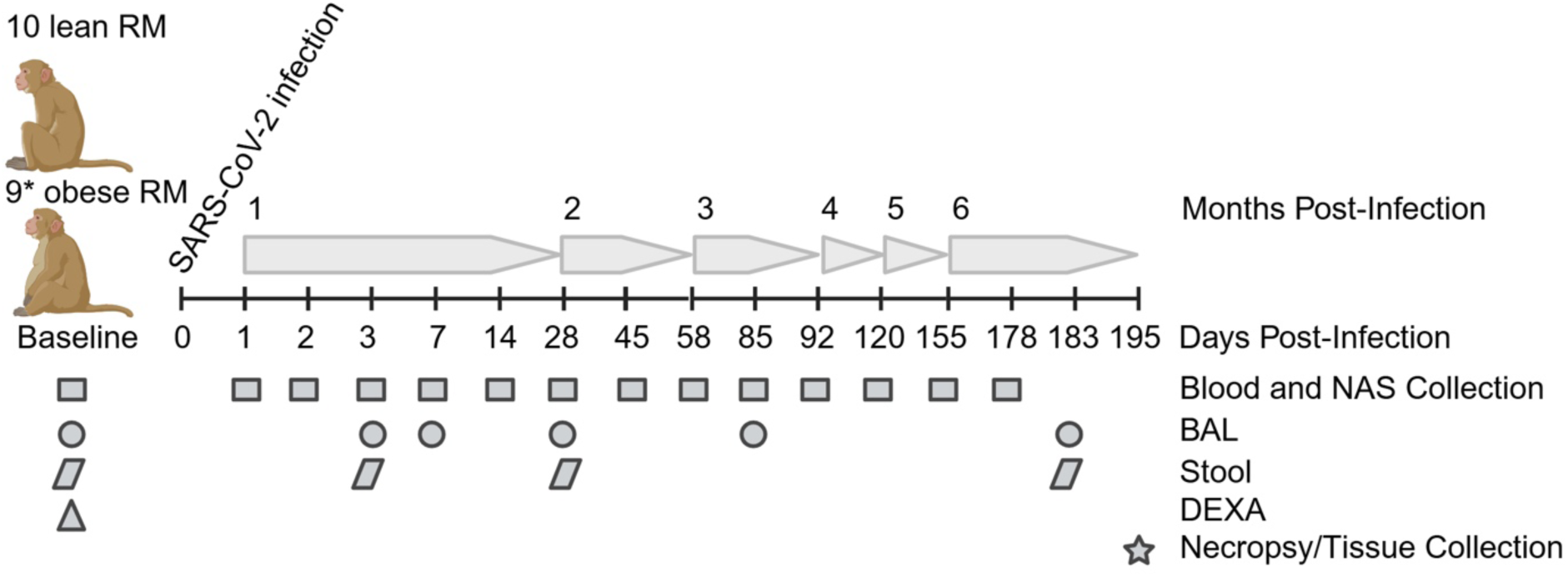
Study design and experimental timepoints. Study timeline and schedule of various procedures and sample collection times are shown. Baseline (B in all figures) samples were taken immediately prior to infection on day 0 for viral dynamics in NAS and BAL samples and at day −26 for the data in Table 1 (including DEXA scans), serology, immune cell and cytokine profiling, and adipokine measurements. When procedures spanned more than 1 day, an average day PI is shown for simplicity. *One obese animal was excluded from the study after developing frank diabetes.

Both groups exhibited similar rapid increases in total viral RNA (vRNA) present in nasopharyngeal swab (NAS) samples that peaked between day 2 and 7 PI (Figure 2A). NAS viral load declined in both groups thereafter; however, both groups still had detectable vRNA at day 85 PI, while vRNA levels in both groups were below the limit of detection (LOD) by day 120 PI. There were no differences in peak viral load (Figure 2B) or vRNA area under the curve (AUC) (Figure 2C). There was a significant positive correlation between vRNA AUC and age at infection (Figure 2D). In both groups, the levels of infectious virus (focus-forming units; FFU) peaked between day 2 and 3 PI, remained elevated through day 7 PI, and declined to below the LOD at day 14 PI (Figure 2E). There were no differences between the groups in peak viral load (Figure 2F) or vRNA AUC (Figure 2G). Overall, the levels of infectious virus in NAS were ∼1000- fold lower than copies of vRNA and declined more rapidly than the levels of vRNA, a pattern similar to that reported in a recent SARS-CoV-2 human challenge study (68).

**Figure 2.**
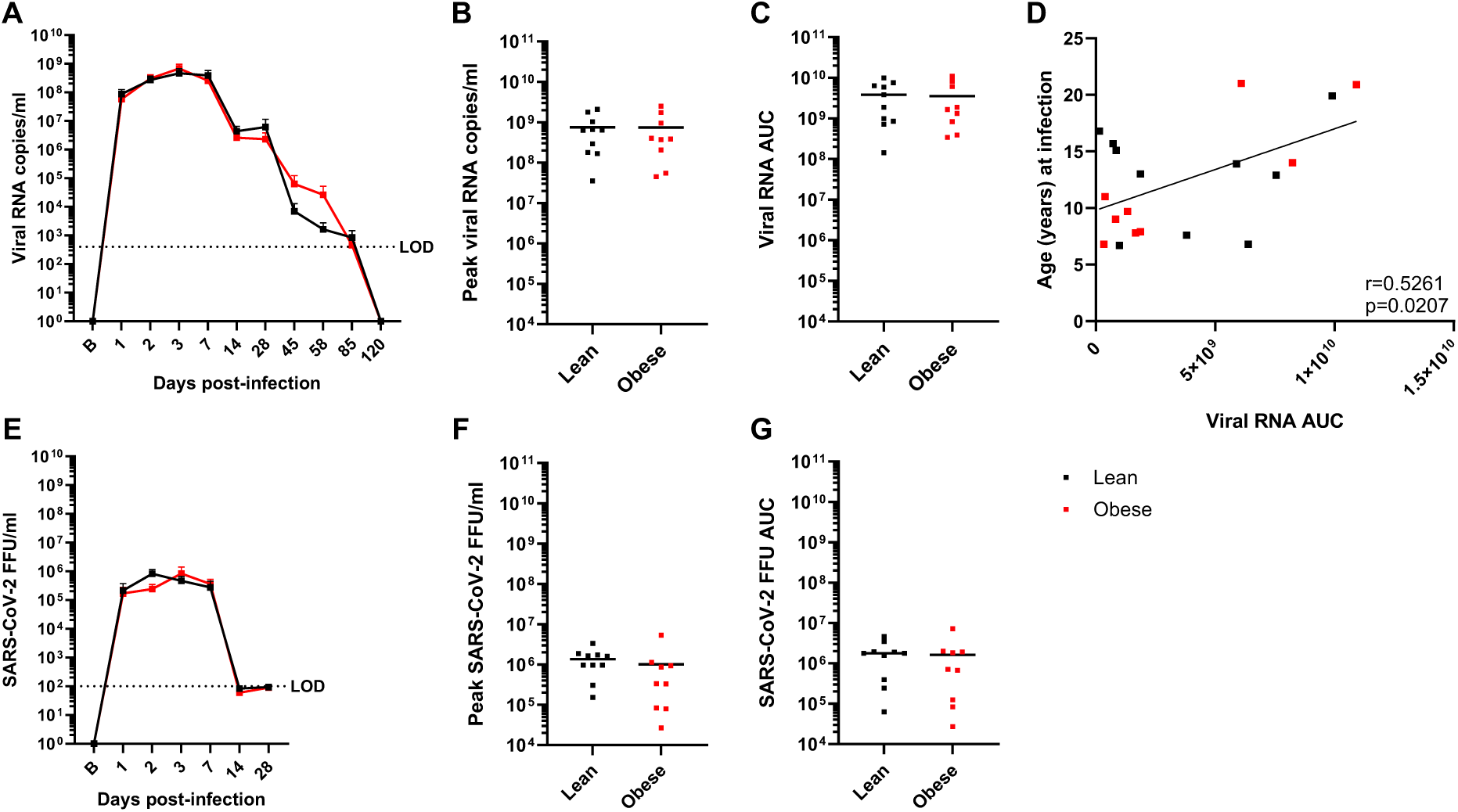
Viral dynamics in NAS samples. A-C. vRNA levels, peak vRNA levels, and vRNA AUC, respectively, in lean and obese groups. D. Correlation between age and vRNA AUC determined by Pearson’s correlation coefficient. E-G. Infectious virus (focus-forming units; FFU) levels, peak FFU levels, and FFU AUC, respectively, in lean and obese groups. Data in panels A and E are means ± SEM. LOD: Limit of detection.

Similar analyses were done with bronchoalveolar lavage (BAL) samples, although fewer time points were obtained due to the invasive nature of BAL fluid collection. In both groups, vRNA levels peaked at day 3 PI and declined after day 7 PI to below the LOD by day 85 PI (Figure 3A). There were no differences between the groups with respect to peak vRNA titer (Figure 3B) or vRNA AUC (Figure 3C), although the obese group trended higher in the latter parameter. This was reflected in a significant positive correlation between vRNA AUC and baseline fat mass (Figure 3D). Infectious virus (FFU) levels in the lean group peaked at day 1 PI and in the obese group at day 7 PI, declining in both groups to the LOD by day 28 PI (Figure 3E), with no difference between the groups in peak viral load (Figure 3F) or vRNA AUC (Figure 3G). Similar to what was observed in NAS, the levels of infectious virus in BAL were ∼10,000- fold lower than copies of vRNA and the level of FFUs declined faster than total vRNA levels; however, the overall levels of vRNA copies and FFUs were more that 10-fold lower in BAL than in NAS. In summary, obesity was associated with greater BAL but not NAS vRNA AUC.

**Figure 3.**
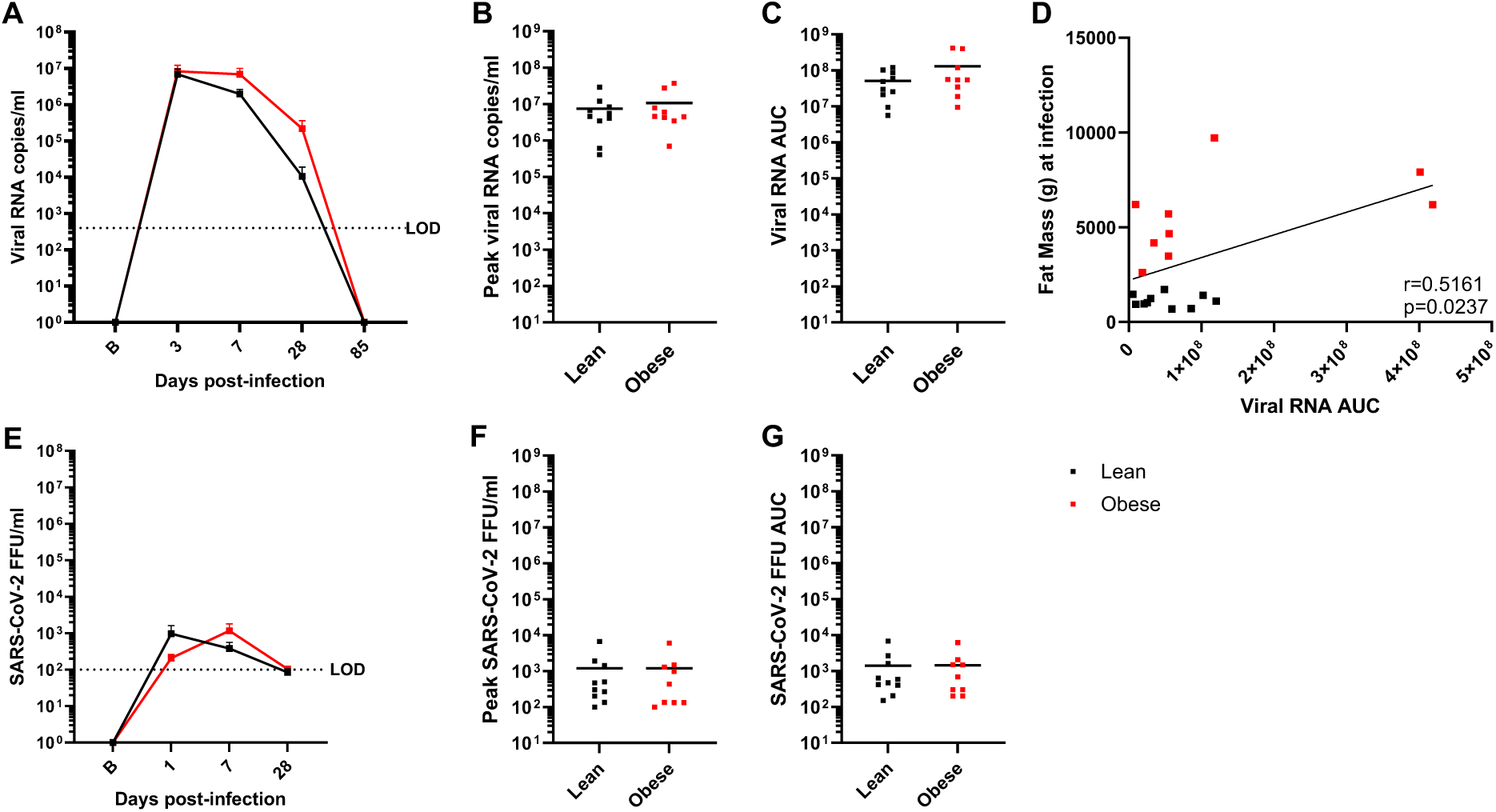
Viral dynamics in BAL samples. A-C. vRNA levels, peak vRNA levels, and vRNA AUC, respectively, in lean and obese groups. D. Correlation between baseline fat mass and vRNA AUC determined by Pearson’s correlation coefficient. E-G. Infectious virus FFU levels, peak FFU levels, and FFU AUC, respectively, in lean and obese groups. Data in panels A and E are means ± SEM. LOD: Limit of detection.

### vRNA persists in multiple locations following acute infection

Longitudinal plasma samples were also tested for vRNA. A single animal tested positive at day 2 PI. All other plasma samples had no detectable vRNA. No detectable vRNA was present in longitudinal cerebrospinal fluid samples (CSF) at day 14 and 30 PI, in 3-month PI WAT biopsies, or 6-month PI (necropsy) cerebellum, pancreas, or lung (left and right caudal, middle, and proximal lobe) samples. Interestingly, 4 lean and 4 obese animals had detectable levels of vRNA in nasal turbinates and a separate lean animal had detectable vRNA in the soft palate at 6 months PI. vRNA in fecal samples was evaluated by RT-PCR using primers that targeted SARS-CoV-2 genomic, antigenomic, and subgenomic vRNA (N2) or only antigenomic and genomic vRNA (NSP14) over the course of infection and recovery. As shown in Figure 4, both lean and obese animals had detectable N2 (Figure 4A) and NSP14 (Figure 4B) vRNA sequences throughout the experimental time course at levels that were higher than three SDs above the mean of the 5 (3 lean and 2 obese) pre-infection samples. There were no obvious differences in the proportions of lean and obese animals that were positive for vRNA at 6 months PI. However, only a few animals had vRNA levels at the 6-month timepoint that were several-fold greater than the background baseline value. The significance of the vRNA levels that were closer to the background level is unclear.

**Figure 4.**
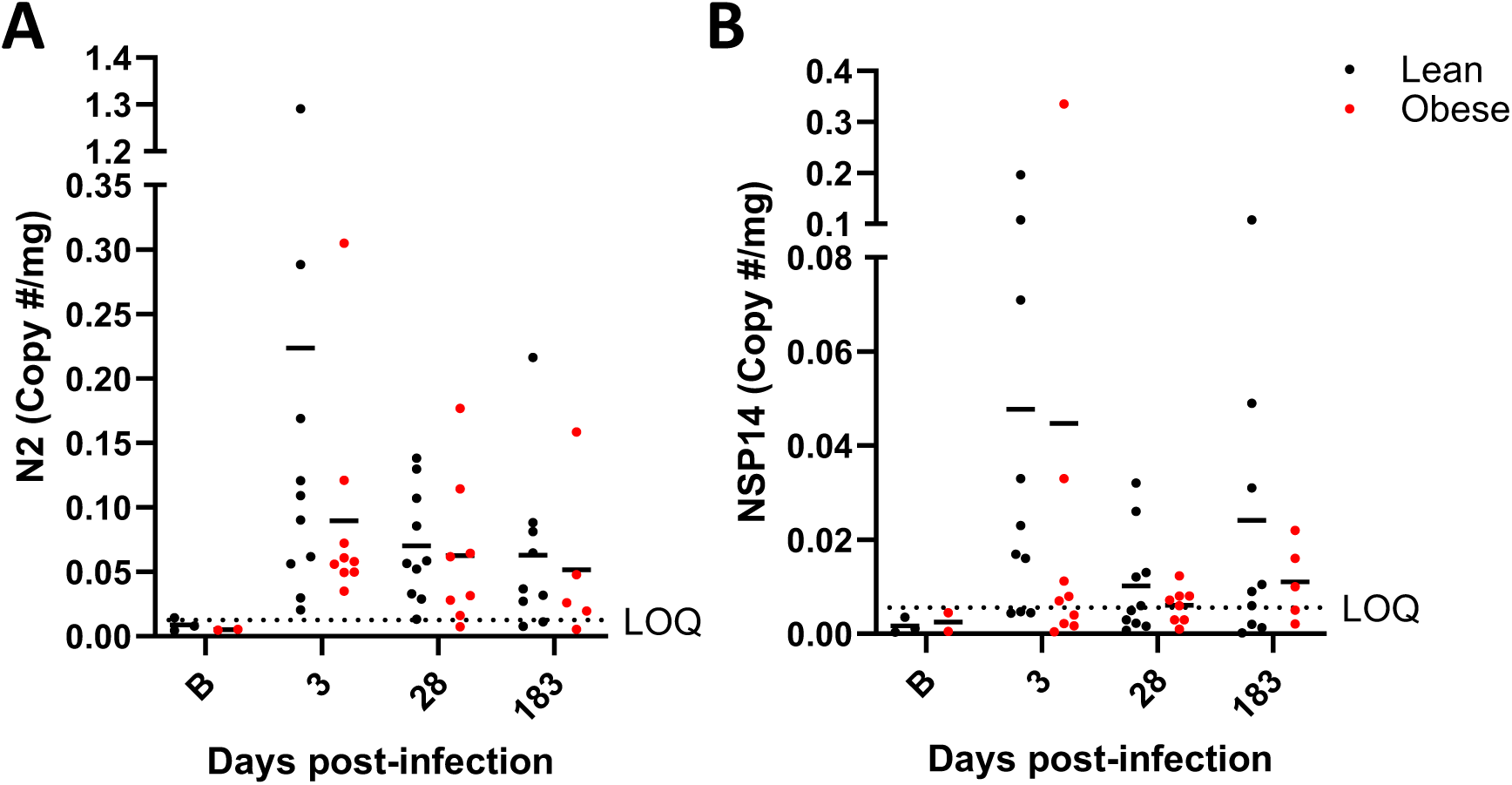
vRNA levels in fecal samples. N2 primer-specific (A) and NSP14 primer-specific (B) vRNA copies/mg in longitudinal stool samples in lean and obese groups. The N2 primer set targets SARS-CoV-2 genomic, antigenomic, and subgenomic vRNA sequences, while the NSF14 primer set targets only antigenomic and genomic vRNA sequences. Horizontal dotted lines represent the limit of quantitation (LOQ) based on 3 SDs above the mean of the values of pre-infection samples (B timepoint on X-axis).

### SARS-CoV-2-specific Ab responses differ in lean and obese animals

Virus-specific Ab responses were measured longitudinally for up to 180 days after SARS-CoV-2 infection. Antiviral IgG responses to the SARS-CoV2 receptor-binding domain (RBD) were measured by ELISA, with Ab titers peaking between day 14 and 28 PI in both groups (Supplemental figure 1A). Lean animals exhibited geometric mean titers (GMT) of 22,000 ELISA Units (EU) by day 14 PI, and this titer remained steady through d 28 PI (GMT=26,000 EU) before declining sharply to 8,300 EU by day 58 PI and eventually falling to 5,400 EU by day 178 PI. Similarly, obese animals mounted antiviral IgG responses of 23,000 EU and 23,500 EU at d 14 and 28 PI, respectively, before declining to 6,800 EU by d 58 PI and later to 4,500 EU by day 178 PI. Clinical studies performed early in the COVID-19 pandemic indicated that SARS-CoV-2 RBD-specific Ab responses declined with an estimated 2.4-m half-life when measured longitudinally during the first 6 months after infection (69). In comparison, RBD-specific Ab responses in lean animals declined with a 2.8-m half-life (95%CI; 2.2-3.8 m) from the peak IgG responses observed at day 28 to day 178. Similarly, Ab responses of obese animals declined with a 2.4-month half-life (95%CI; 2.1-2.9 m), which was not significantly different from the lean group (p=0.33 by unpaired t test). Together, these results indicate that the antiviral Ab responses observed in SARS-CoV-2-infected macaques did not differ between the groups but were similar to those observed after natural human SARS-CoV-2 infection.

Analysis of neutralizing Ab titers (representing a combination of antiviral IgG and IgM responses) showed a somewhat more variable and prolonged plateau compared to RBD-specific IgG responses (Supplemental figure 1B). The 50% focus-reduction neutralizing titer (FRNT50) observed after infection of lean animals reached a GMT of 170 FRNT50 at both day 14 and day 28 PI, rising slightly to between 320-330 FRNT50 at d 58 to d 85 PI, before declining to 26 FRNT50 by day 178 PI. Obese animals mounted similar neutralizing responses with GMTs of 103-110 FRNT50 at days 14-28 PI, dipping to 69 FRNT50 at d 58 PI before returning to 160 FRNT50 at day 85 PI and then eventually declining to a GMT of 35 FRNT50 by day 178 PI.

In a separate analysis of the Ab decay rate kinetics from day 28 to day 178 PI, the FRNT50 of the lean group declined with a 1.8-m half-life (95%CI: 1.4-2.6 m), whereas Ab responses in the obese group declined with a 4.1-m half-life (95%CI: 2.2-35 m), a statistically significant difference (p=0.04 by unpaired t test). The 6.5-fold decline in FRNT50 in the lean group from day 28 to day 178 PI is similar to the 7-fold decline in neutralizing titers observed from 2 to 14 months after infection among clinical cases of COVID-19 (69), in contrast to the 3.1-fold decline in FRNT50 seen in the obese group. Thus, the neutralizing Ab levels achieved in obese animals trended lower than in the lean group and decayed at a significantly slower rate. Overall, RBD-specific IgG responses were similar in both groups, while neutralizing RBD-specific Ab dynamics differed between lean and obese animals.

### SARS-CoV-2 infection exerts differential effects on lung pathology in lean and obese animals

As described in the Methods section below, three specific lesions (bronchioalveolar hyperplasia, lymphocytic interstitial pneumonia, and perivascular lymphoid aggregates) were selected as the parameters for a lung pathology scoring system (Supplemental table 1). These histologic features were evaluated for each lung lobe and each animal was assigned a composite score. All animals exhibited some degree of interstitial and perivascular lymphocytic infiltration. Bronchoalveolar hyperplasia, characterized by clusters of hyperplastic epithelial cells reflecting earlier damage at the level of the terminal bronchiole, alveolar duct, and alveolus, was observed in 8 of 19 animals (4 in the lean group and 4 in the obese group). Nodular aggregates of perivascular lymphocytes were observed in six animals (3 obese and 3 lean). Additional microscopic lesions not included in the scoring system were present in one or more animals. They were considered related to SARS-COV-2 infection and included organized thrombi, prior alveolar hemorrhage, alveolar fibrin accumulation, granulomatous inflammation centered on hemoglobin crystals, type II pneumocyte hyperplasia, bronchiolar hyperplasia, interstitial fibrosis, and luminal mucus. Representative images of the scored features are shown in Supplemental figure 2.

Figure 5 illustrates and contrasts the gross and microscopic findings of representative lungs with low and high pathology scores. The lungs with a low score (Figure 5A) had a pathology score of 7 on microscopic examination. The lung lobes were grossly and histologically unremarkable (Figure 5C,E). The left lung lobes and right caudal lung lobes of the animal that had a high score (Figure 5B) were mottled pink and dark red and were soft rather than normally spongy on palpation at necropsy. Microscopically, the pathology score was 29 due to the presence of moderate bronchoalveolar hyperplasia and chronic interstitial pneumonia (Figure 5D,F).

**Figure 5.**
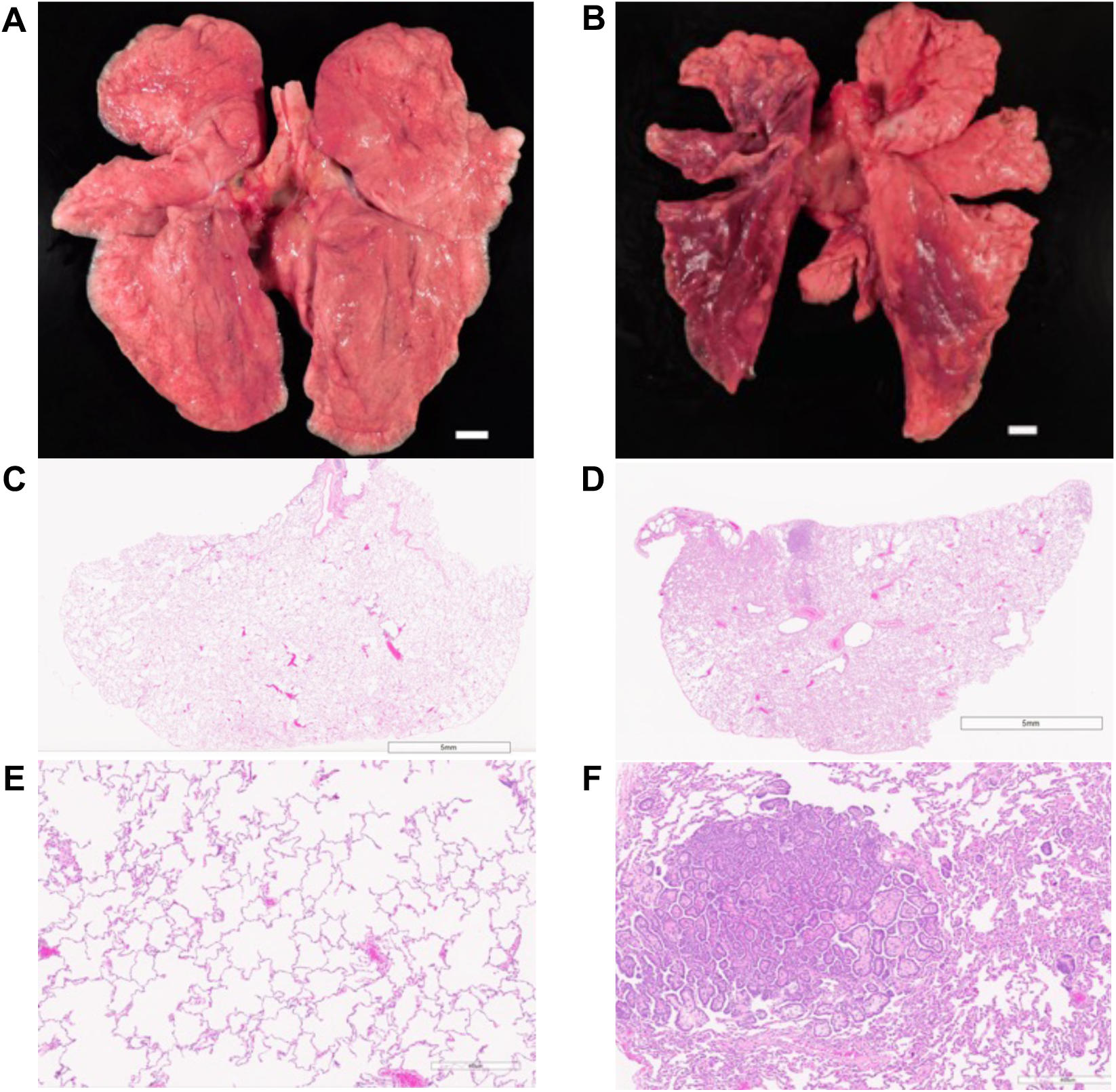
Gross and microscopic pulmonary findings of representative animals with low and high pathology scores. Necropsy photographs of the lung lobes and attached trachea of lungs with low (A) and high (B) lung pathology scores. H&E-stained micrographs at the subgross level (C,D) and at a higher magnification (E,F) of the left caudal lobe appear below each animal’s gross photograph. Lung lobes with low score were grossly and histologically unremarkable (A,C,E). All the left lung lobes and the right caudal lung lobe with high score exhibited discoloration and were mottled pink and dark red (B). Moderate bronchioloalveolar hyperplasia was present and characterized by clusters of proliferative epithelial cells surrounding a fibrovascular core (D,F). Bars in A and B are 1 cm.

As shown in Figure 6A, composite scores ranged from 4 to 29, with the higher scores representing more severe lesions. Five of six animals with higher scores (14 to 29) were in the obese group. Two of seven animals with lower scores (5 and 7) were also in the obese group. The fat mass in the obese group ranged from 2608 to 9715 g (20.94 to 48.03 % body fat) and the pathology scores ranged from 5 to 29. The fat mass in the lean group ranged from 690 to 1719 g (7.59 to 16.89 % body fat) and the pathology score ranged from 4 to 22. The average pathology score trended higher in the obese group but did not reach statistical significance (p=0.0972). However, individual pathology scores were significantly associated with baseline fat mass (Figure 6B). Individual pathology scores were also significantly associated with vRNA AUC in both NAS (Figure 6C) and BAL (Figure 6D) samples. Individual pathology scores also trended to correlate with FFU AUC in NAS (p=0.0504) but not in BAL. Individual pathology scores also trended higher with age, but did not reach statistical significance (p=0.0642). Overall, increased lung pathology at necropsy was associated with higher fat mass and acute vRNA AUC in NAS and BAL.

**Figure 6.**
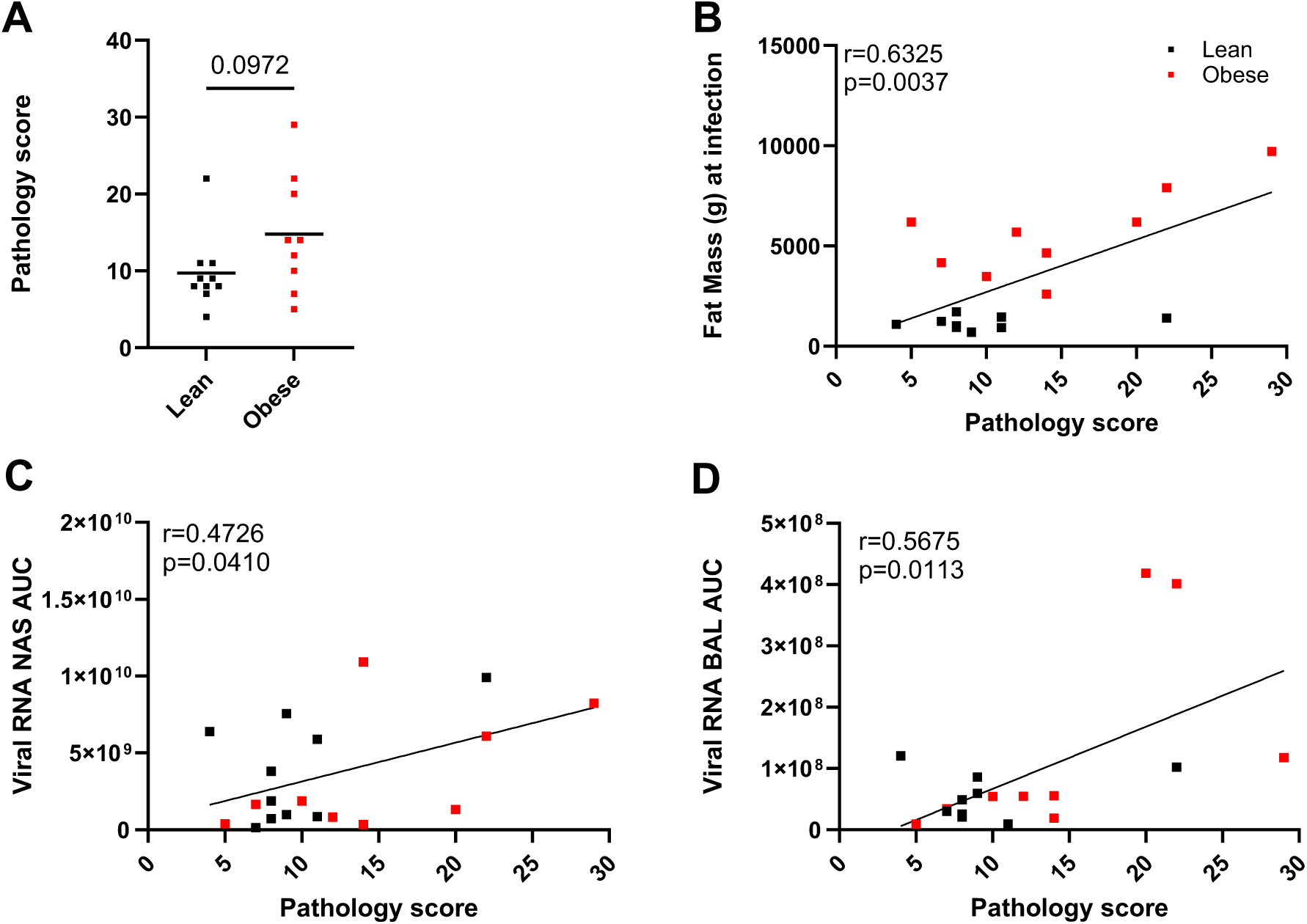
Quantitation of lung pathology. A. Pathology scores in lean and obese animals differentiated by unpaired 2-tailed t test. Correlation of pathology scores with baseline fat mass (B), vRNA AUC in NAS (C) and BAL (D) determined by Pearson’s correlation coefficient.

### SARS-Cov-2 infection differentially affects blood immune cell profiles in lean and obese animals

To assess the overall effects of SARS-CoV-2 infection on the cellular components of the innate and adaptive arms of the immune system, longitudinal changes in blood immune cell populations were assessed by flow cytometry. CD14 and CD16 staining was used to investigate monocyte populations and CD169 was used for assessment of global monocyte activation. As shown in Supplemental figure 3A, the levels of intermediate, classical, and CD169+monocytes each differed at baseline between the groups (p=0.0238, p=0.0324, and p=0.0201, respectively). The levels of non-classical monocytes varied slightly over the time course and the pattern significantly differed between lean and obese animals. The levels of more proinflammatory intermediate monocytes peaked at day 3 PI in both groups (p<0.0001 vs B) (Supplemental figure 3B), with a second peak at day 28 occurring only in the lean group (p=0.0055 vs Obese). Thus, the levels of intermediate monocytes also varied over time following infection, and the response differed between lean and obese animals. The levels of classical monocytes varied over time following infection and differed between the groups (Supplemental figure 3C). The most striking effect of SARS-CoV-2 infection was the massive increase in CD169 expression on CD14+ monocytes in both groups on days 2-3 PI (p<0.0001 vs B) that quickly resolved by day 14 PI (Supplemental figure 3D). Thus, the response of all categories of monocytes to infection differed in lean and obese animals.

We also examined CD16+ NK cells, important for cytotoxic responses, and CD56+ NK cells, involved in proinflammatory responses, as well as proliferation of NK cell subsets as assessed by Ki67 staining. There were no differences in any of the categories between groups at baseline. The levels of CD16+ and CD56+ NK cells were significantly affected by infection and the pattern of response differed by group (Supplemental figure 4A,B), including an increased level of CD56+ NK cells at d 3 PI only in lean animals (p=0.0295 vs obese). The levels of Ki67 activation in CD16+ and CD56+NK cells were also affected by infection (Supplemental figure 4C,D) and the pattern differed between groups, although both groups exhibited a peak in Ki67+CD16+ NK cells at d 7 PI (lean p=0.0008 vs B and obese p=0.0004 vs B).

Changes in circulating T cell profiles over the course of SARS-Cov-2 infection and recovery are shown in Supplemental figure 5. There were differences in the baseline proportions of CD69+CD4+, CD69+CD8+, CD38CD4+, and CD38CD8+ T cells (p=0.0408, p=0.0015, p<0.0001, and p=0.0311, respectively). The frequencies of total CD4+ and CD8+ cells over time were affected similarly by infection in both groups. (Supplemental figure 5A,B). We also assessed longitudinal changes in the frequencies of the T cell activation/proliferation markers CD69, Ki67, and CD38 (Supplemental figure 5C-H). In each case, levels varied following infection and the pattern differed between the groups, although both groups exhibited peaks at day 2 PI in CD69+CD4+ (lean p=0.0002 vs B and obese p=0.0005 vs B) and CD69+CD8+ T cells (lean p=0.0003 vs B and obese p=<0.0001 vs B) and at day 3 PI in CD38+CD4+ T cells (lean p=0.0011 vs B and obese p=<0.0001 vs B). CD38+CD8+ T cells exhibited a peak at day 3 PI only in the obese group (p=<0.0001 vs B).

We also monitored CD4+ and CD8+ naïve (Supplemental figure 6A,B), central memory (Tcm) (Supplemental Figure 4C,D), and effector memory (Tem) (Supplemental figure 6E,F) T cell subsets during SARS-Cov-2 infection. All categories were affected by infection, and the pattern was different in lean and obese animals, except in CD4+ Tem cells, where longitudinal changes were similar in both groups (Supplemental Figure 4E).

To determine if SARS-CoV-2 infection affected the acute humoral immune response, we longitudinally monitored B cells as well as B cell proliferation and changes in the frequencies of memory B cell subsets. The frequencies of CD3-CD8-CD79a+ B cells (Supplemental figure 7A), Ki67+ B cells (Supplemental figure 7B), and naïve, IgM memory, or switched B memory cells (Supplemental figure 7C-E) all varied following infection, but the responses of lean and obese animals were different in each case.

Overall, SARS-CoV-2 infection altered the frequencies of almost all the immune cell categories analyzed, and most of these exhibited responses that differed between lean and obese animals. However, while significant fluctuations in immune populations were observed over the study time course, particularly during acute infection, no persistent differences in immune cell populations between the groups were seen at 6 months, even in cases where the groups differed significantly at baseline.

### SARS-CoV-2 infection alters BAL and plasma cytokine profiles

Cytokine profiling of longitudinal samples of BAL fluid and plasma was performed to evaluate altered cytokine levels potentially associated with SARS-CoV-2-induced inflammation and to determine the presence of any persistent or delayed changes. BAL samples from 10 lean and 9 obese animals were analyzed at baseline and days 3, 7, 28, 72, and 180 PI. Plasma samples from a subset of 3 lean and 3 obese animals were also analyzed in the same assay format at baseline and days 3, 28, and 180 PI. Analytes that exhibited significant changes over time are shown in Figure 7. The levels of CXCL10/IP-10 (Figure 7A) and IL-15 (Figure 7B) in BAL fluid peaked in both the lean and obese groups during the acute phase of infection, with CXCL1-/IP-10 levels exhibiting the most dramatic increase. Levels of CXCL10/IP-10 and IL-15 returned to baseline levels by day 28 PI. The acute increases in CXCL10/IP-10 and IL-15 seen in BAL fluid were also seen in plasma (Figure 7C,D), and plasma PD-L1 also exhibited a transient increase in both groups at day 3 PI (Figure 7E). In contrast, increases in several plasma analytes, including BDNF, CCL2/MCP-1, CCL5/RANTED, CCL11/Eotaxin, IL-8/CXCL8, and TNF-a were only seen at day 180 PI (Figure 7F-K). The other analytes assayed either did not exhibit longitudinal changes or were present at levels below the limit of quantitation (LOQ) of the assay.

**Figure 7.**
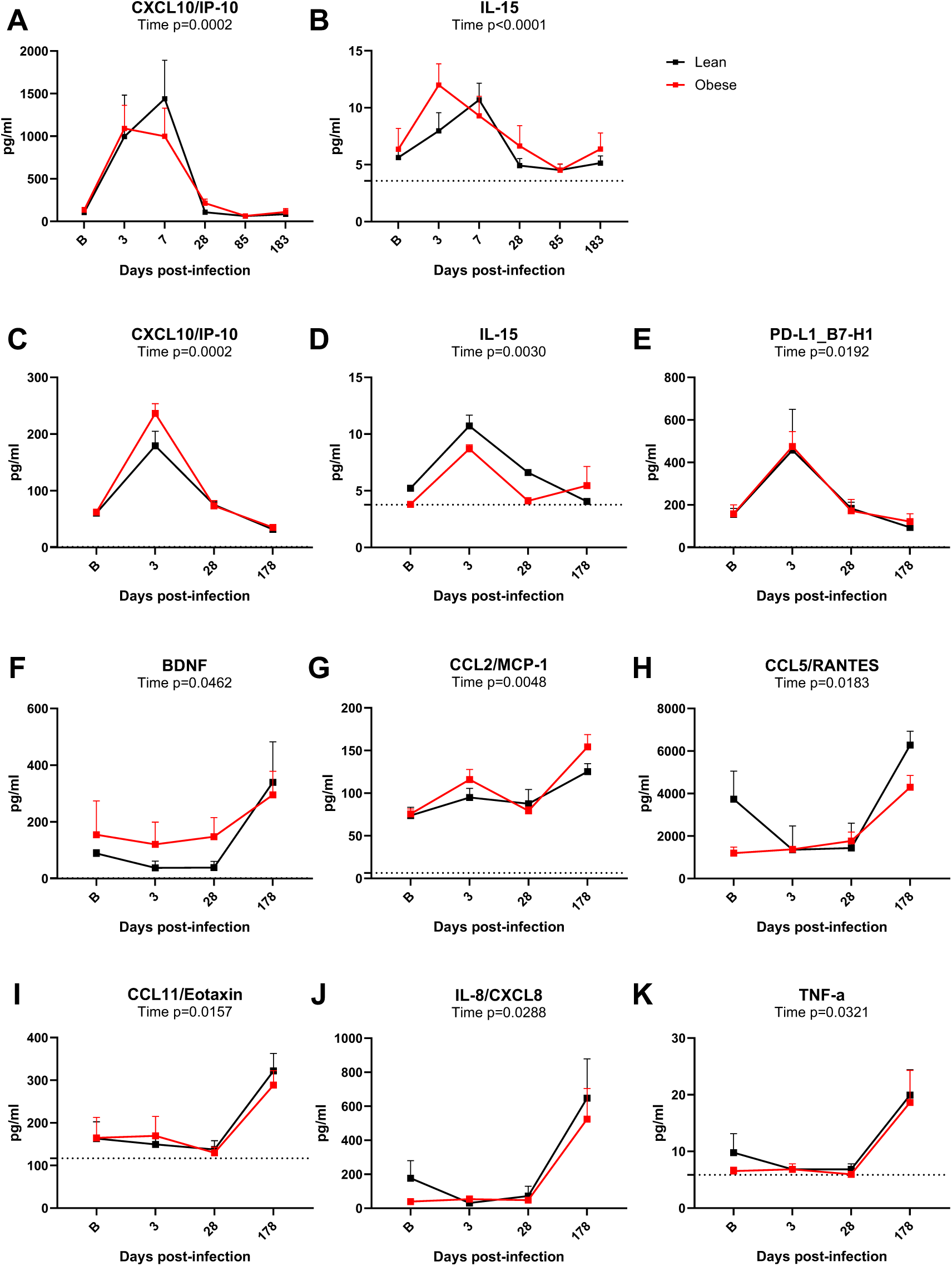
BAL and plasma cytokine profiles. Longitudinal changes in BAL CXCL10/IP-10 (A) and IL-15 (B) in BAL from lean and obese animals determined with a Luminex multiplex assay. Longitudinal changes in plasma CXCL10/IP-10 (C), IL-15 (D), PD-L1 (E), BNDF (F), CCL2/MCP-1 (G), CCL5/RANTES (H), CCL11/Eotaxin (I), IL-8/CXCL-8 (J), and TNF-α (K) analyzed using the same assay. All data are means ± SEM. Significance determined using mixed-effect analysis with Dunnett’s post-hoc for multiple comparisons test. Horizontal dotted lines in panels B, D, G, I, and K depict limit of quantitation (LOQ). In panels A, C, E, F, H, and J, the LOQ was too close to 0 to show.

### SARS-Cov-2 infection differentially affects BW in lean and obese animals

As shown in Figure 8A, BW changes in response to infection differed between the lean and obese groups. Both lean and obese animals lost similar amounts of their baseline weight (−4.2% lean and −4.5% obese) by day 7 PI. However, the lean group then rebounded, while the obese group continued to lose additional weight up until day 28 PI (−7.9%). The lean group recovered their initial BW around day 28 PI, but the obese group did not fully rebound until approximately day 85 PI. The average BW of animals in the lean group continued to increase throughout the subsequent time course, resulting in an overall 16.3% weight gain after 6 months that was significantly greater (p=0.0022 versus baseline) than the 5.8% overall increase observed in the obese group (p=NS versus baseline). Figure 8B shows the absolute BW of the lean and obese groups at baseline and at day178 PI. As indicated, the overall weight gain in the lean group at 6 months PI was significant, whereas that in the obese group was not. In summary, both groups exhibited an initial BW loss, but the lean group rebounded more quickly and was significantly heavier versus baseline at the end of the study, while the obese group took much longer to rebound, and BW was not significantly different from baseline at study end.

**Figure 8.**
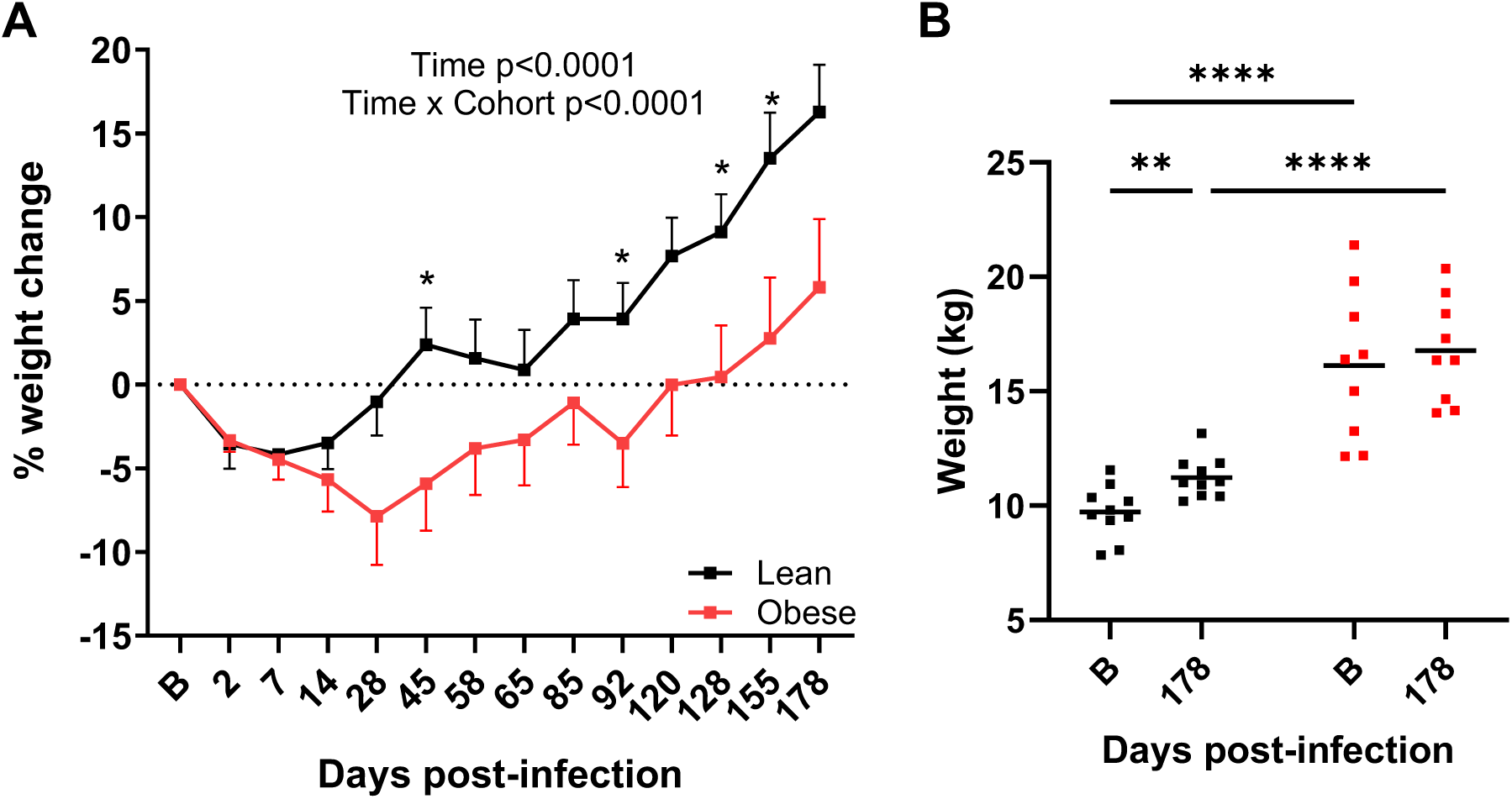
Body weight changes after SARS-CoV-2 infection. A. Longitudinal changes in body weight in lean and obese animals. Significance determined using mixed-effect analysis with Dunnett’s post-hoc for multiple comparisons test. Data are mean ± SEM. B. Body weight changes between baseline and 6 months PI. Significance determined by 2-way repeated-measures ANOVA with post-hoc pair-wise comparisons. *, p<0.05; **, p<0.01; ****, p<0.0001.

### SARS-CoV-2 infection affects long-term metabolic status in both lean and obese animals

We measured FBG and FBI levels to determine the effects of SARS-CoV-2 infection on glucose control and insulin sensitivity. As shown in Figure 9A, FBG levels in both groups were similar at baseline but declined in the lean group over the study time course (p=0.015). As shown in Figure 9B and Table 1, FBI levels were significantly higher in the obese group at baseline (p<0.05) through day 28 PI but then significantly declined by d 58 PI (p=0.0204). As described above, the higher FBI in the obese group was reflected in the similar FBG levels in the two groups due to compensatory hyperinsulinemia in the obese animals. However, FBI levels decreased in the obese group following the decrease in BW shown in Figure 8A. Thus, SARS-CoV-2-induced BW loss in the obese group improved insulin sensitivity as measured by HOMA-IR, which persisted through initial rebound in BW (Figure 9C), driven mainly by the decrease in FBI in the obese group. Although the obese group began to gain BW at day 155 PI, this was not sufficient to restore the baseline FBI. Baseline HbA1c levels also differed between the groups and changed in parallel over the subsequent study time course (Figure 9D).

**Figure 9.**
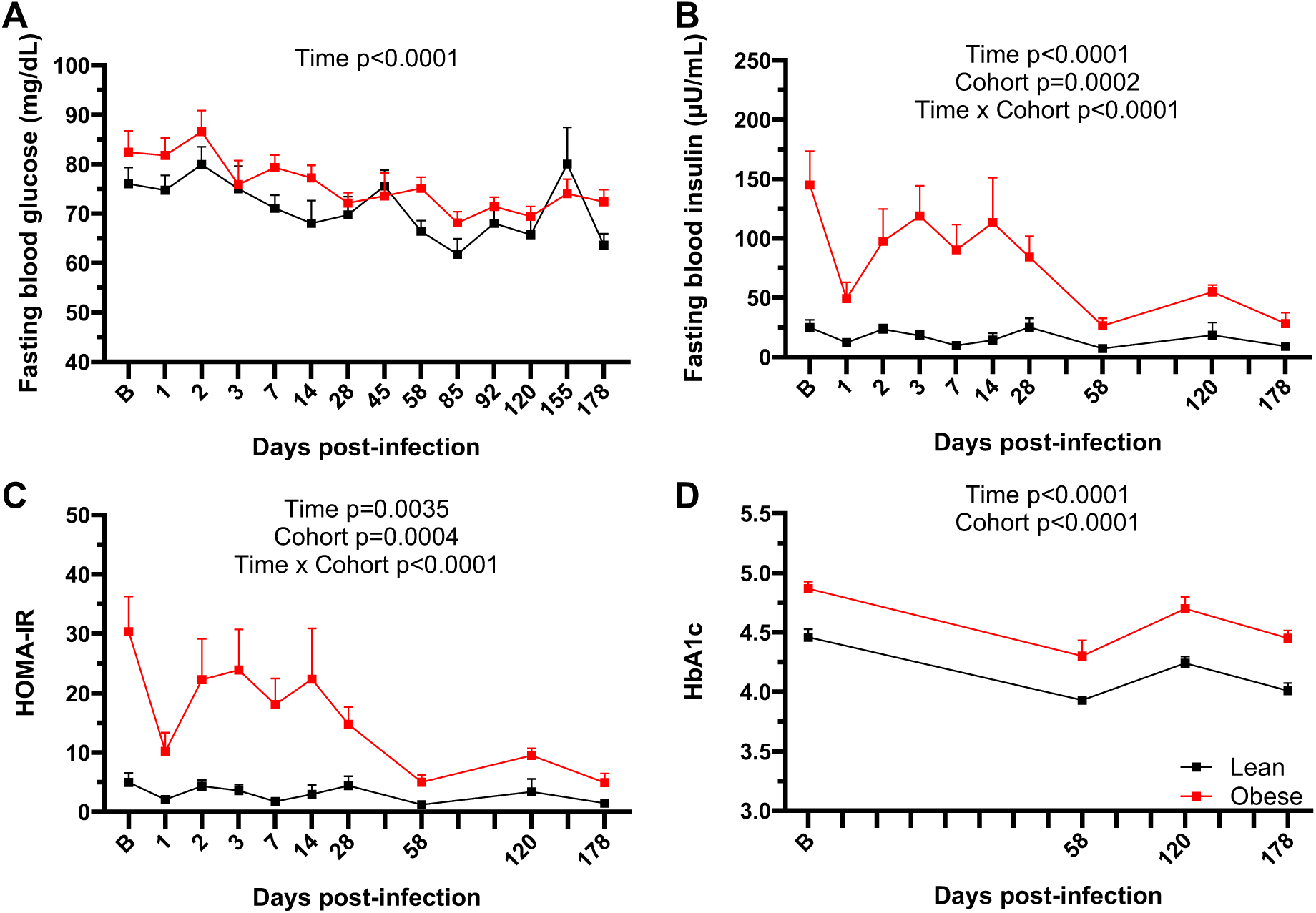
Glucose homeostasis during SARS-CoV-2 infection. FBG (A), FBI (B), and HOMA- IR (C) levels and HbA1c values (D) in lean and obese animals over the study time course. All data are means ± SEM. Significance determined using mixed-effect analysis with Dunnett’s post-hoc for multiple comparisons test.file.

### Baseline metabolic status influences specific adipokine responses to SARS-CoV-2 infection

Longitudinal changes in the circulating levels of the major WAT-derived cytokines adiponectin and leptin were determined by ELISA and RIA, respectively. As shown in Figure 10A, plasma adiponectin levels at baseline were significantly higher in the lean group compared to the obese group (p=0.0337). The lean group exhibited a modest decrease in circulating adiponectin at day 3 that rebounded by day 14 PI, but that then decreased dramatically by ayd 58 PI to a level that was maintained through day 180 PI (p<0.0001). Adiponectin levels in the obese group were unchanged through day 28 PI and then significantly declined by d 58 PI (p=0.0076). By day 58 PI, the levels of the lean group had declined to levels not significantly different from those in the obese group. Plasma leptin levels in the lean group were significantly lower than those in the obese group at baseline (p<0.0001) and remained constant (Figure 10B). Leptin levels in the obese group declined slightly by day 14 PI, and those levels were maintained through day 180 PI. (Figure 10B). The adiponectin/leptin ratio (ALR) is an important biomarker of inflammation and cardiometabolic risk (45–47). The ALR in the lean group at baseline was 4-fold higher than that in the obese group (p=0.0019), but declined to a value not significantly different from that exhibited at baseline by the obese group (p=0.393; Figure 10C,D). These data demonstrate that a greater decrease in adiponectin in the lean group drove the significant decrease in the ALR to a level similar to that exhibited by the obese group.

**Figure 10.**
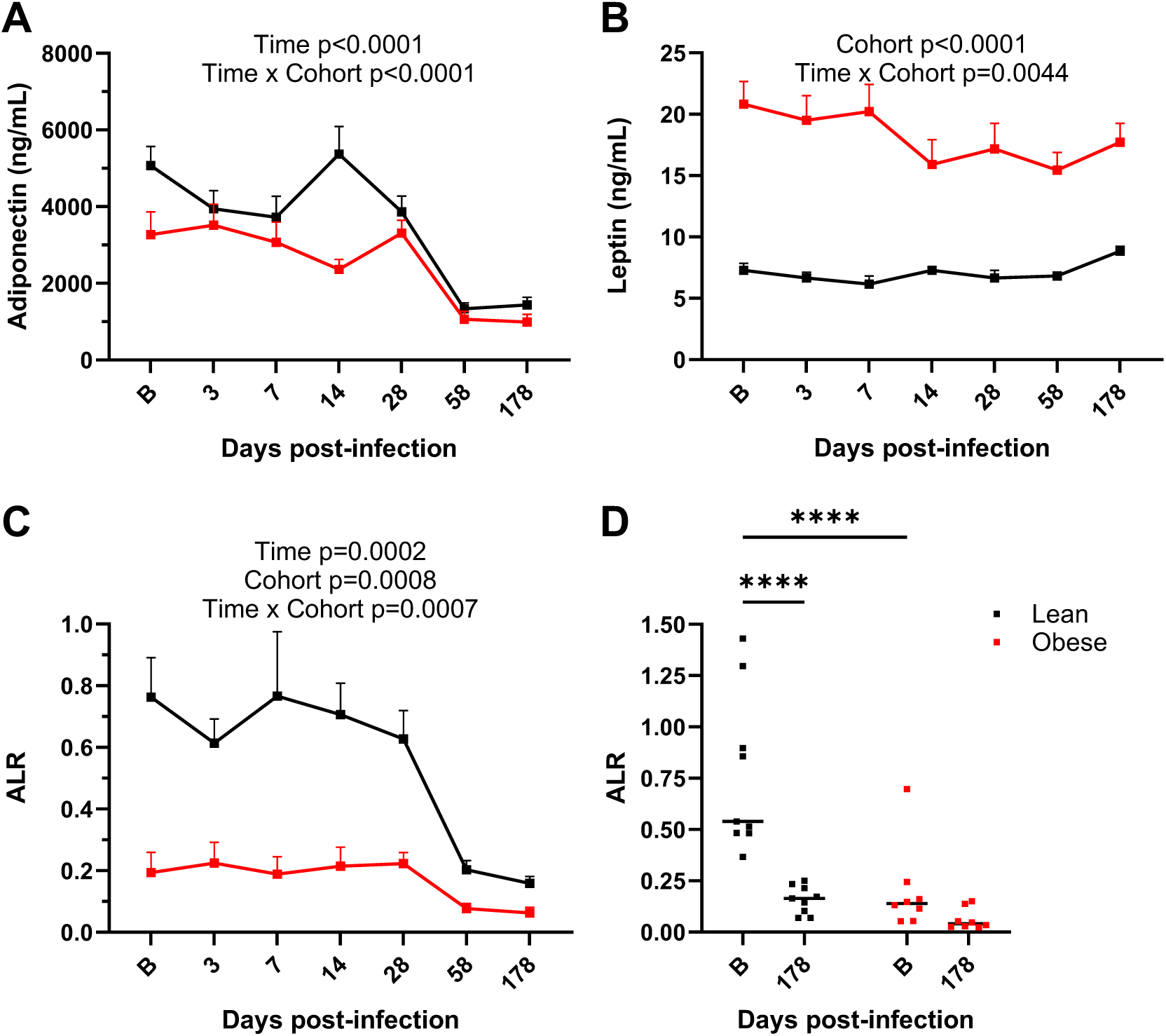
Plasma adiponectin and leptin levels. A. Total plasma adiponectin levels in lean and obese groups determined by ELISA. B. Plasma leptin levels in lean and obese groups determined by RIA. C. Adiponectin/leptin ratio (ALR) in lean and obese groups. D. ALR in lean and obese groups at baseline and d 180 PI. Data in panels A-C are means ± SEM. Significance determined using mixed-effect analysis with Dunnett’s post-hoc for multiple comparisons test. Significance in panel D determined by 2-way repeated measures ANOVA with post-hoc pair-wise comparisons. ****, p<0.0001.

### SARS-CoV-2 infection exerts differential effects on BT and daytime/nighttime activity in lean and obese animals

Star-Oddi loggers were implanted in a subset of 5 lean and 5 obese animals and programed to record BT and physical activity measurements each h for the duration of the study time course. As shown in Supplemental figure 8, the lean and obese animals in this subset were similar in age but had significantly different BW, reflecting the differences in the overall experimental cohort. Supplemental figure 9A shows the circadian rhythm of BT with peak temperatures occurring during daylight h and dropping during nighttime from a window of time just prior to infection through day 14 PI. At baseline, the range of BT was greater in lean animals (max 37.5 ^0^C vs min 34.6 ^0^C = range of 2.1 ^0^C) when compared to obese animals (max 37.1 ^0^C vs min 35.6 ^0^C = range of 1.5 ^0^C). This larger range in the lean group was due to a lower nighttime BT. Obese animals exhibited an increased daytime peak of BT at day 3 PI (green box in Supplemental figure 9A). To determine if there were long-term effects on BT, we examined a daytime window (11 AM-1 PM) at baseline, day 3, and day 185 PI (Supplemental figure 9B). Both groups had a transient rise in daytime BT at day 3 PI that was significant compared to baseline only in the lean group. Next, we examined BT during a nighttime window (1-4 AM) at baseline and day 3 and dat 185 PI (Supplemental figure 9C). Obese animals had a significantly higher nighttime BT at baseline compared to lean animals. Obese animals had a transient rise in nighttime BT at day 3 PI that was significant compared to the lean group. Interestingly, BT at day 185 PI in the lean group was significantly higher than at baseline or day 3 PI. Therefore, the larger range of daytime vs nighttime BT in the lean group was reduced by SARS-CoV-2 infection.

To determine if SARS-Cov-2 infection had a long-term effect on physical activity levels, we examined defined windows (11 AM-1 PM for daytime activity and 1-4 AM for nighttime activity) throughout the study time course. Daytime averages for the lean cohort ranged from 20 to 150 and for the obese cohort from 8 to 35 throughout the study, demonstrating that, overall, lean animals were more active than obese animals, but that daytime activity levels were not affected by SARS-CoV-2 infection. Figure 11A shows the pattern of nighttime activity (1-4 AM) in the lean and obese groups, and Figure 11B shows average nighttime activity at baseline and at 6 m PI (average of days 185-188 PI). Nighttime activity, indicating restlessness, significantly increased in both cohorts, but to a greater extent in the lean cohort (3-fold).

**Figure 11.**
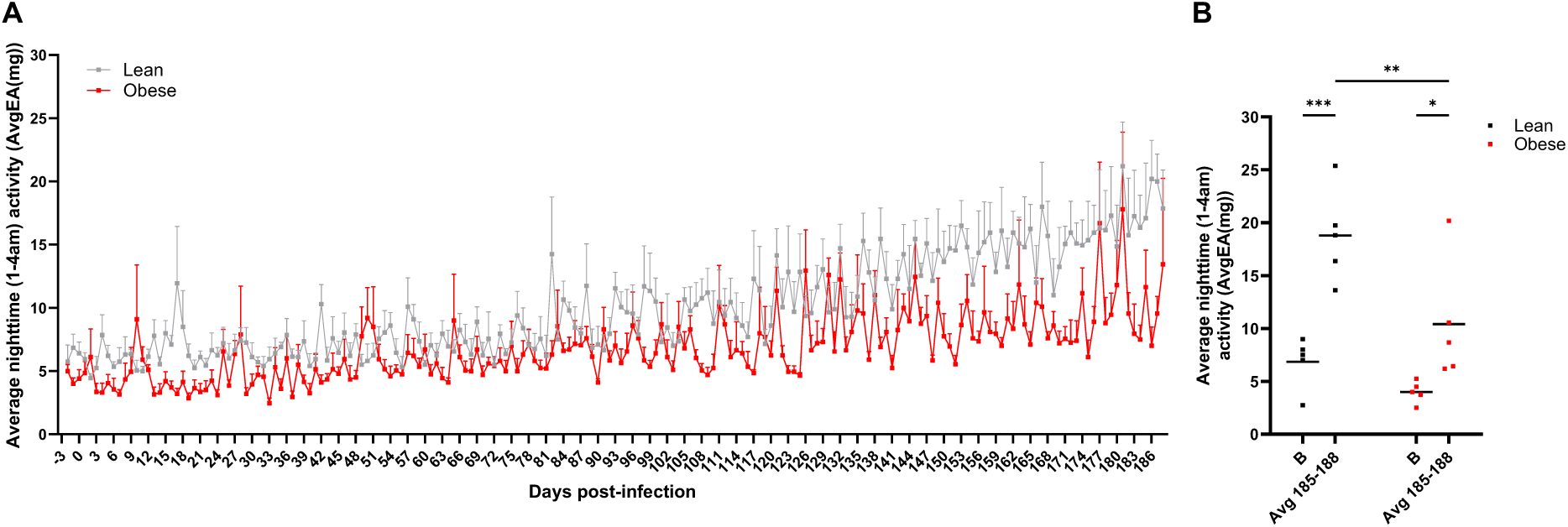
Nocturnal activity following SARS-CoV-2 infection in lean and obese animals. A. Nighttime activity (1-4 AM) in lean and obese animals from d −3 (baseline) to 186 PI. B. Average nighttime activity (1-4 AM) at baseline (d −3, B) and 185-188 PI. Data in panel A are means ± SEM. External acceleration (EA) units reflect any movement greater than standard gravity detected by the data logger implant. Significance determined by 2-way repeated-measures ANOVA with post-hoc pair-wise comparisons. *, p<0.05; **, p<0.01; ***, p<0.001.

## Discussion

This study utilized a rhesus macaque model of SARS-CoV-2 infection to ascertain the effects of pre-existing obesity and reduced insulin sensitivity on the exacerbation of ongoing metabolic dysfunction in obese animals and/or the development of new metabolic disease in lean animals. We also monitored an array of non-metabolic parameters to determine the potential influence of prior metabolic disease on other physiological processes. We were particularly interested in any differential responses between lean and obese animals to SARS-CoV-2 infection and which responses in either group developed or persisted following the acute phase of infection, with the latter potentially comprising aspects of PASC.

Almost every parameter assessed in this study exhibited changes following SARS-CoV- 2 infection, and the majority differed between lean and obese animals. As discussed in more detail below, some parameters changed more robustly in obese animals, while others changed more robustly in lean animals. Thus, persistent effects of SARS-CoV-2 infection are both obesity-dependent and independent.

Viral dynamics in NAS and BAL were similar in both groups, with infectious virus levels decaying more rapidly than total vRNA levels in both cases. However, a greater vRNA AUC was correlated with older age in NAS and with individual baseline BW in BAL, even though the average vRNA only trended higher in the obese group. This association of increased fat mass with BAL vRNA AUC but not FFU AUC may explain some of the association of obesity with adverse outcomes in COVID-19, since multiple SARS-CoV-2 gene products that may be expressed by persistent noninfectious virus have been shown to exert cytotoxic effects in vitro (70, 71).

The presence of persistent vRNA in nasal turbinates, soft palate, and stool (the latter presumably reflecting persistent infection of GI tissues) following an initial mild acute disease course, combined with other persistent phenotypic changes, supports the SARS-COV-2- infected rhesus macaque as a valid preclinical model of PASC. Human studies have also reported persistent vRNA in stool samples, although at lower frequencies (72, 73), and that likely reflect viral persistence in the GI tract (74). These data suggest that SARS-CoV-2-infected macaques also harbor viral reservoirs that may contribute to components of PASC (75) (76).

The adaptive immune response to SARS-CoV-2 infection reflected in total SARS-CoV-2 RBD-specific IgG titer was rapid and, after a modest drop at 2 months PI, was sustained thereafter. The levels of anti-RBD IgG/M neutralizing Ab, on the other hand, increased less in the obese group during acute infection and decreased more slowly. Thus, pre-existing obesity appeared to specifically affect neutralizing Ab decay rates.

Baseline obesity also affected lung pathology assessed at 6 m PI. The lung pathologies employed for scoring were chosen because they represented a chronic response to injury likely due to prior SARS-CoV-2 infection; however, none of them were considered specific to SARS CoV-2 infection. The significant association of vRNA AUC in NAS and BAL with individual pathology scores provides strong support for the validity of the scoring system. All observed lung pathologies were generally minimal to mild, which is consistent with the minimal/limited clinical disease observed in these animals. All animals had at least minimal scores for lymphocytic interstitial and perivascular infiltrates in one or more lung lobes. This is consistent with background findings in adult and aged animals in the rhesus macaque colony from which the study animals were obtained (ADL and LAMC, pers. comm). Individual animals tended to score higher in one category than others. While the average lung pathology score only trended higher in the obese group, baseline fat mass in individual animals was associated with a higher pathology score, suggesting that pre-existing obesity increases the risk for long-term lung dysfunction. Another important aspect of our findings is the presence of significant lung pathology in the absence of overt clinical symptoms, similar to what we have observed previously with the delta variant (57), suggesting that SARS-CoV-2 infection can exert persistent effects on lung function even in the absence of a severe clinical presentation.

Studies of chronic lung pathology in humans are largely limited to the evaluation of patients with severe terminal COVID-19 disease (77, 78). There are no published studies of post-acute lung pathology in SARS-CoV-2-infected NHPs. The longest disease course previously studied in rhesus macaques was 36-42 d PI and used the beta variant (79). The lung pathology seen in the current study mirrors that seen in Boszormenyi et al., but many of these animals had additional lesions, specifically nodular perivascular lymphocytic aggregates and bronchioloalveolar hyperplasia. This may reflect a difference in virus strain since, in our previous studies, the delta variant also produced more lung pathology during acute infection (57). Our findings of post-acute lung pathology are consistent with a recent report of persistent lung repair in convalescent COVID-19 patients (80).

SARS-CoV-2 infection resulted in changes in the frequency of multiple circulating immune cell populations, including various categories of monocytes, T cells, NK cells, and B cells, and most of these responses differed between lean and obese animals. As described below, however, the differential response of immune cell profiles in lean and obese animals was not associated with differential responses in BAL of plasma cytokines.

Cytokine profiling of BAL fluid and plasma revealed that there were minimal longitudinal changes evident in BAL, while more analytes were altered in plasma, with the latter changes mainly occurring at the end of the study period. This was somewhat unexpected, since we had assumed that acute changes at least would be more robust in BAL. The modest response in BAL may reflect the mild nature of infection with the delta variant, while the delayed changes seen in several plasma analytes suggests that the levels of specific cytokines may be persistently altered. Of the plasma analytes that exhibited increases at the 6-m timepoint, BNDF (81), CCL2/MCP-1 and TNF-a (82), CCL5/RANTES (83), and CCL11/Eotaxin (84) have been previously reported to be increased in long COVID patients. The fact that many analytes were below the limit of quantitation in the Luminex assay employed suggests that a more sensitive and comprehensive analysis may be warranted, especially for plasma analytes in light of the small sample size of our plasma analysis.

The lean and obese groups also differed in the effect of SARS-CoV-2 infection on BW, where, despite an initial loss in both groups, lean animals rebounded quickly and gained significant BW by 6 m PI, while obese animals exhibited a more prolonged initial weight loss and lagged the lean group in recovering baseline BW. FBG levels trended downward in both groups during the study, while FBI levels preferentially decreased in the obese group, presumably linked to the decreased BW, such that insulin sensitivity in this group was improved. Overall, SARS-CoV-2 infection induced a greater loss of BW in obese animals, which improved their insulin sensitivity, an effect which persisted even after this group regained their baseline BW. These data suggest that SARS-CoV-2-induced weight loss may temporarily reduce obesity-associated insulin resistance. Our findings differ from those reported by Palmer et al. in SARS-CoV-2-infected African green monkeys (85), in which non-obese animals developed persistent hyperglycemia after SARS-CoV-2 infection. Potential reasons for this discrepancy include the difference in species, our use of the delta variant vs the earlier alpha variant employed by Palmer et al., and/or potential sex differences, since our study involved only males, while the comparable animals in the study of Palmer et al. were 90% female.

Another important finding of this study was the progressive decline in circulating adiponectin in conjunction with a minimal change in circulating leptin, such that the ALR in the lean group was decreased to the level characteristic of the obese group by 4 m PI. This finding suggests that SARS-CoV-2 infection increases long-term cardiometabolic risk in lean animals but does not further increase existing cardiometabolic risk in obese animals. Previous human studies have reported the association of low adiponectin levels (41, 42) or a decreased ALR (44–46) with severe acute COVID-19 disease and mortality, but ours is the first study, to our knowledge, that analyzed the time course of adiponectin and leptin levels after the acute phase of infection. The observed preferential decline in adiponectin in the lean group is strikingly similar to the decline in adiponectin and the ALR in lean but not obese animals following SIV infection and prolonged antiretroviral therapy (ART) that we reported previously (86). In that study, the effects on the ALR could be due to either persistent effects of SIV or represent a comorbidity of ART. If the former is the case, it is tempting to speculate that long-term effects on cardiometabolic risk may be shared between SARS-CoV-2 and SIV.

The changes in BW described above in both groups likely reflect, at least in part, changes in fat mass with possible decreases in adipocyte size and/or number which could potentially decrease adiponectin levels, since WAT is a major source of plasma adiponectin (87). However, since circulating adiponectin levels continued to decline, particularly in the lean group, even after both groups began to regain weight, circulating adiponectin levels do not appear to be associated with WAT mass. Thus, some other aspect of SARS-CoV-2 infection must be the basis for the decrease in circulating adiponectin and, as a result, the ALR. Since a decrease in BW principally reflects decreased subcutaneous and visceral WAT depots, we cannot exclude the possibility that SARS-CoV-2 infection-associated changes in bone marrow WAT contribute to the observed changes in plasma adiponectin, since bone marrow WAT is also a potential source of circulating adiponectin (88). It is also notable that circulating adiponectin levels declined at the same time that insulin sensitivity increased; thus, the discordance between adiponectin levels and insulin sensitivity suggests altered metabolic homeostasis following recovery from acute SARS-CoV-2 infection.

Our initial hypothesis was that obesity could contribute to more severe acute infection and the development of some aspects of PASC through a greater number of WAT cells subject to SARS-CoV-2 infection as well as altered adipocytokine profiles. Our data do not provide any direct evidence for the first mechanism, since no vRNA was detectable in WAT at the earliest biopsy taken at 3 m, although virus could have been present during acute infection. On the other hand, the previously described relationship between a decreased ALR and WAT dysfunction (47, 48) and the selective decrease in the ALR observed in lean animals does support a relationship between SARS-CoV-2 infection and subsequent long-term changes in WAT function. These results suggest that additional studies, including changes in WAT architecture and cell profiles following acute SARS-CoV-2 infection, are warranted.

In addition to the long-term decrease in the ALR that was unique to the lean group, two other parameters that were persistently altered preferentially in lean vs obese animals were BT and activity. Specifically, average daytime BT was increased during acute infection and decreased to below baseline levels only in the obese group, while average nighttime BT only increased in lean animals at 6 m PI but average nighttime BT in obese animals was significantly higher at baseline and trended higher during acute infection but returned to baseline levels by 6 m PI. Thus, a persistent decrease in daytime BT was only seen in obese animals, while a persistent increase in nighttime BT was only seen in lean animals. Alterations in BT have been recently reported in long COVID patients (89).

Persistent increases in nocturnal activity were also noted following SARS-CoV-2 infection. Lean animals exhibited a significant increase in nighttime activity that was particularly evident towards the end of the study period, while obese animals were somewhat less active at baseline and exhibited a smaller increase in nocturnal activity by the end of the study period.

These findings are consistent with recent reports of insomnia and sleep disturbances in long COVID (90, 91). The development of specific aspects of PASC in lean animals, including those associated with metabolic dysfunction such as a decreased ALR, and increased nighttime activity, suggests that, while preexisting obesity can increase the risk for some components of PASC such as long-term lung pathology, other aspects of PASC can develop in the absence of pre-infection obesity.

A major result of this study was the identification of multiple parameters, including activity levels and specific cytokine levels, that were consistently altered in the majority of convalescent animals months after the resolution of acute infection and viremia. This high incidence of PASC based on quantifiable parameters in a well-controlled preclinical model of mild-to-moderate disease is much greater than estimates of PASC prevalence in the human population (5, 92). This discrepancy may result from the reliance of the latter on self-reporting of symptoms and the fact that many of the parameters we measured, while reflecting pathology, may not necessarily cause symptoms that would be apparent to an affected individual.

We acknowledge that this study does have several limitations. For logistical and budgetary reasons, it was not possible to include females, although future studies using females are clearly warranted based on the reported sex differences in multiple components of PASC (93–95). Indeed, the scope and magnitude of the effects we report here may well be greater in females in light of the greater extent of PASC in women. For the same reasons, we did not include an uninfected control group; however, we feel that the longitudinal study design, in which each animal serves as its own control, precluded the need for an uninfected control group. Another limitation was the inability to perform certain assessments that would have been informative due to limitations on the feasibility of certain assays and procedures in the ABSL-3 barrier facility that animals were housed in for the entirety of the study. A final limitation is the use of obese animals whose weight gain was induced by an obesogenic WSD. We consider differences between the lean and obese groups to be mainly due to obesity and/or associated insulin resistance, but we cannot exclude the possibility that the differences seen reflect, at least in part, an interaction between SARS-CoV-2 infection and WSD that is independent of obesity per se.

The strengths of this study include the use of phenotypically well-characterized experimental groups, identical pre-infection SARS-CoV-2-naïve status, and the unique opportunity to collect longitudinal samples for an extended period and to procure multiple tissue samples at necropsy. Together, these features make this the most comprehensive pre-clinical study of PASC features to date that also includes types of data unobtainable in clinical studies. Notably, our findings of persistent phenotypic changes following initial mild-to-moderate disease and the presence of detectable vRNA in several tissue sites at 6 m PI establishes the SARS-CoV-2-infected rhesus macaque model as a valid model for PASC and suggests that the true prevalence of PASC in the human population may be significantly underestimated.

## Methods

### Sex as a biological variable

Our study exclusively examined male rhesus macaques. It is unknown whether the findings are relevant for female macaques.

### Animals

This study employed 10 lean, metabolically healthy and 9 obese, insulin-resistant adult male rhesus macaques (one animal in the original obese group of 10 animals was excluded due to the development of frank diabetes part way through the study). Baseline demographics of the two groups are shown in Table 1 and discussed above. Study animals were housed individually on a 12-hour light-dark cycle in a climate-controlled, Animal Biosafety Level 3 (ABSL-3) facility that that allowed protected interaction with other rhesus macaques in the same room. Lean animals were fed a control diet containing 15% of calories derived from fat (Purina 5000/5052; LabDiet, Gray Summit, MO), while obese animals were fed an obesogenic WSD containing 36% of calories derived from fat (Purina 5LOP, LabDiet). Animals in the obese group had been maintained on this diet for at least a year. Sedations for procedures were induced with ketamine HCl (5-15 mg/kg) (Covetrus, Dublin, OH) or a Tiletamine HCl and Zolazepam HCL mixture (3-5 mg/kg) (Telazol™; Zoetis Inc., Kalamazoo, MI), unless otherwise noted.

### Body composition

Determination of baseline total, lean, and fat mass, as well as bone mass and BMD were assessed using dual-energy X-ray absorptiometry scanning (Lunar iDXA, GE HealthCare, Chicago, IL). After an overnight fast, animals were sedated with 3-5 mg/kg Telazol and positioned in a prone position during a standard scan.

### SARS-CoV-2 infection

After completion of baseline assessments, all animals were infected via intratracheal and intranasal routes with 1 × 10^6^ pfu of the SARS-CoV-2 delta variant (isolate hCoV-19/USA/MND-HP05647/2021 B.1.617.2; delta variant, WCCM; Cat. #NR-56116; lot 70047614, BEI Resources) as previously described (57).

### Nasal swab (NAS) collection

NAS were collected using a Copan FLOQSwab nasal swab (Cat. #23-600-952, Thermo Fisher Scientific, Waltham, MA) that was inserted into the nares, gently rotated, and then immersed in 0.4 ml of PBS containing 1x protease inhibitor cocktail (Cat. #78415, Thermo Fisher Scientific) or Dulbecco’s Modified Eagle’s Medium (DMEM) containing 1x penicillin–streptomycin-glutamine (Cat. #10378016, Thermo Fisher Scientific) and 1x Antibiotics-Antimycotic solution (Cat. #15240062, Thermo Fisher Scientific).

### Scopeless BAL collection

Animals were anesthetized, intubated, and positioned in right lateral recumbency. A sterile 8 to 10-Fr catheter was passed through the endotracheal tube and into the trachea. The catheter was advanced until resistance was felt. BAL was performed by infusing three to four 10-ml aliquots of normal saline followed by immediate aspiration of the saline infusate after each 10-ml infusion.

### SARS-CoV-2 RNA (vRNA) levels and infectious virus titers

Blood, NAS, BAL fluid, CSF, and tissue samples were processed for vRNA extraction and viral load and infectious virus titers (focus-forming units; FFU) were determined as previously described (57).

### vRNA detection in fecal samples

Prior to purification of total RNA from stool, approximately 1.5 g of frozen stool was added to a pre-weighed Zymo Research DNA/RNA Shield Fecal Collection Tube (Cat. #R1137, Zymo Research, Irvine, CA) to determine individual sample mass. Samples were then thawed on ice and homogenized by vortexing for approximately 30 seconds or until a homogenous solution was formed. Sample solution was then clarified via centrifugation at 4000 X g for 20 minutes at 4 °C. Supernatant was subsequently removed and filtered through a 22-μM syringe. 250 μL of sample filtrate was then added to 750 μL of Trizol LS reagent (Cat. #10296010, Thermo Fischer Scientific) and total RNA was extracted with a Zymo Research RNA Clean & Concentrator-25 kit (Cat. #R1018, Zymo Research). cDNA was synthesized via reverse transcription-PCR on 1 μg of total RNA, deoxynucleotide triphosphates, and Random Primers with Moloney Murine Leukemia Virus Reverse Transcriptase (Cat. #28025013, Invitrogen, Waltham, MA). Quantitative PCR was then carried out on each cDNA product using SYBR Green (Invitrogen) and SARS-CoV-2-specific primers targeted to SARS-CoV-2 genomic, antigenomic, and subgenomic vRNA sequences (N2) or only antigenomic and genomic vRNA sequences (NSP14). N2 primers were TTACAAACATTGGCCGCAAA (forward) and GCGCGACATTCCGAAGAA (reverse) and NSP14 primers were TGGGGYTTTACRGGTAACCT (forward) and AACRCGCTTAACAAAGCACTC (reverse). cDNA synthesis was also performed on synthetic SARS-CoV-2 RNA of known copy number, which served as a standard curve to determine the SARS-CoV-2 copy number. The SARS-CoV-2 copy number detected in stool samples was normalized to the initial input weight of stool in mg.

The data shown in Fig. 4 were derived from pre-infection samples from 3 lean and 2 obese animals to determine the background level of signal, from all 19 animals (10 lean and 9 obese) at days 3 and 28 PI, and from 14 animals (9 lean and 5 obese) that had samples available at day 180 PI. The average number of copies/mg from the pre-infection samples was determined and all subsequent timepoints with values greater that 3 SDs above the mean of the pre-infection samples were considered to represent persistent vRNA.

### SARS-CoV-2-specific IgG levels

Plasma samples were tested for SARS-CoV-2 receptor-binding domain (RBD)-specific IgG Abs by ELISA. 96-well ELISA plates (Cat. #CLS9018, Millipore Sigma, Burlington, MA) were coated with 100 μL of recombinant SARS-CoV-2 RBD T478K protein (Cat. #40592-V08H91, Sino Biological, Wayne, PA) at a concentration of 1 μg/mL in phosphate-buffered saline (PBS), incubated overnight at 4°C and stored frozen at −20°C until use. Plates were thawed at room temperature (RT), the coating antigen was removed by flicking the plates over a sink and gently tapping on paper towels, blocked for 1 h at RT with 5% nonfat dry milk (Kroger brand) in PBS containing 0.05% Tween (dilution buffer) and washed once with PBST (wash buffer). Heat-inactivated plasma samples were serially 3-fold diluted in 100 μL dilution buffer in the ELISA plates. Plates were incubated at RT for 1 h, followed by the addition of 50 μL of 10% hydrogen peroxide and further incubated for 30 minutes at RT. The plates were washed 3 times with wash buffer and 100 μL of 1:2000 dilution of horseradish peroxidase (HRP)-tagged anti-monkey IgG detection Ab (Cat. #GAMon/IgG(Fc)/PO, Nordic MUbio, Susteren, the Netherlands) was added. Plates were incubated at RT for 1 h and washed 3 times with wash buffer. 100 μL of colorimetric detection reagent containing 0.4 mg/mL of o-phenylenediamine and 0.01% hydrogen peroxide in 0.05 M citrate buffer (pH 5) was added, and the reaction was stopped after 20 minutes by the addition of 100 μL of 1M HCl. OD at 490 nm was measured using a VersaMax ELISA plate reader (Molecular Devices, LLC, San Jose, CA). An internal positive control serum standard was included on all ELISA plates to normalize titers. Ab titers were determined by log-log transformation of the linear portion of the curve using 0.1 OD as the endpoint and performing conversion of the final values. Samples were tested in duplicate and paired samples with >25% coefficient of variation were repeated.

### SARS-CoV-2 neutralizing antibodies

Plasma was diluted 1:10 into DMEM/2% FBS, then serially diluted 1:3 (down to 1:810). An equal volume of diluted plasma was mixed with ∼100 FFUs of virus (final dilutions were 1:20-1:1620) and incubated 1 h at 34°C. Virus/Ab mix was then plated on Vero cells expressing transmembrane protease, serine 2 and human angiotensin-converting enzyme 2 (Vero E6-TMPRSS2-T2A-ACE2; obtained through BEI Resources, NIAID, NIH, NR-54970) and incubated 1 hour and overlayed with carboxymethyl cellulose. After incubation for 24 hours at 34°C, cells were fixed and foci detected as previously described (57). Values below the limit of detection (1:20) were assigned an arbitrary value of one-half of the limit of detection (1:10).

### Necropsies

All animals were euthanized at 6 months PI. Representative tissue samples from the major organs were collected and stored in Trizol for RNA isolation and also fixed in 10% neutral-buffered formalin for at least 7 days, then processed, embedded in paraffin, sectioned at 5 microns and stained with H&E for histological examination.

### Collection and analysis of lung tissues

Lung lobes and trachea were removed en bloc and photographed. Approximately one quarter of each lung lobe was then clamped off from the edge with hemostats; the tissue was removed and used for determination of vRNA. With the hemostats in place, 10% neutral-buffered formalin was slowly infused via the airways in the remainder of each lobe. Once inflated, the main bronchus of each lobe was tied off and immersed individually in jars of formalin for a minimum of 7 days. Representative sections from each lung lobe were then processed, embedded in paraffin, sectioned at 5 microns and stained with H&E. Nine lung slides, which included two slides of the caudal lobes and one each of the other lung lobes, were scanned at 40x with an Aperio AT2 microscope slide scanner (Leica Biosystems, Deer Park, IL). Stained and scanned slides were examined by two board-certified veterinary pathologists (co-authors ADL and LMAC) blinded to animal group assignments. After an initial review, histologic findings were then recorded and aggregated. Those findings considered to represent responses to chronic injury and that occurred in multiple animals were used as the basis of subsequent scoring. Specifically, bronchioalveolar hyperplasia, lymphocytic interstitial pneumonia, and perivascular lymphoid aggregates were selected as being associated with SARS-CoV-2 infection. A score was assigned for each of these three histologic features identified in every lung lobe examined of individual animals and were added together for a final score. Scoring criteria are detailed in Supplemental table 1 and representative images of these lesions are shown in Supplemental figure 2.

### PBMC immune profiling by flow cytometry

Data were acquired on an LSR II flow analyzer (Becton Dickinson, Franklin Lakes, NJ) and analyzed using FlowJo version 10 (TreeStar, Ashland, OR). Antibodies used are listed in Supplemental table 2 and gating strategies are illustrated in Supplemental figure 10.

### BAL and plasma cytokine profiles

Longitudinal BAL samples from all animals were analyzed by the ONPRC Endocrine Technologies Core using the 37-plex NHP XL Cytokine Luminex Performance Premixed Kit (R&D Systems cat #FCSTM21, Minneapolis, MN) on a Luminex 200 platform (R&D Systems). Plasma samples from a subset of lean and obese animals (n=3 each) were analyzed in the same assay.

### Adipokine measurements

Plasma adiponectin was assayed by ELISA (ALPCO total adiponectin cat# 80-ADPHU-E01, Salem, NH), and plasma leptin by RIA (Millipore cat# HL-81K, Billerica, MA), both in the ONPRC Endocrine Technologies Core.

### Data logger implants

A subset of animals (5 lean and 5 obese) was anesthetized and positioned in ventral recumbency followed by sterile prep and draping of the scapular region. A linear 2-cm skin incision was created in the left/right parascapular space, and a subcutaneous pocket was created by blunt dissection. A pre-programmed heart rate and temperature telemetry unit (Star-Oddi DST milli-HRT ACT, Gardabaer, Iceland) was inserted and the defect closed with a subcutaneous stay suture and continuous intradermal 4-0 Monocryl.

### Statistics

Statistical analyses were performed using Prism software. Longitudinal data for parameters that had baseline values were analyzed for effects of SARS-CoV-2 infection per se (Time in figures), intrinsic differences between the lean and obese groups (Cohort in figures), and differences in the longitudinal response to SARS-CoV-2 infection between the lean and obese groups (Time x Cohort in figures) using mixed-effect analysis with Dunnett’s post-hoc for multiple comparisons test. Correlations were determined using Pearson’s correlation coefficient. AUC was calculated starting at T=0. A p value <0.05 was considered significant.

### Study approval

Animals were managed according to the ONPRC animal care program, which is fully accredited by the Association for Assessment and Accreditation of Laboratory Animal Care, International, and based on guidelines, laws, regulations and principles stated in the Animal Welfare Act (United States Department of Agriculture), Guide for the Care and Use of Laboratory animals (Institute for Laboratory Animal Research), and the Public Health Service Policy on Humane Care and Use of Laboratory Animals (National Institutes of Health). Experimental procedures were reviewed and approved by the OHSU Institutional Biosafety Committee and the ONPRC IACUC.

### Data availability

All raw data associated with data presented in the manuscript are included in the Supporting Data Values files.

## Author contributions

PK, JBS, and CTR designed the study. KAS, GMW, LB, CNK, DLT, CZ, CMM, ADL, LMAC, MAK, HB, CP, MH, JH, MS, RCZ, ST, AT, AJH, SRC, MKS, and DNS conducted experiments and acquired data. KAS, GMW, ADL, LMAC, LG, AJH, ML, SRC, SEK, MKS, DNS, JBS, PK, and CTR interpreted results. KAS, MKS, DNS, PK, and CTR wrote the manuscript. MKS, DNS, JBS, PK, and CTR are co-senior authors. All authors reviewed and edited the manuscript.

## Acknowledgements

This study was supported by a research grant from the Polybio Research Foundation to SRC and NIH grants R01 DK122843-05S1 to JBS, PK, and CTR, R01 DK132225 to MKS, DNS, PK, and CTR, and P51 OD011092 for operation of the ONPRC. We acknowledge the assistance of the ONPRC Endocrine Technologies Core for analysis of plasma cytokine and adipokine levels. The Aperio AT2 instrument employed for analysis of lung histology was supported by NIH grant S10 OD25002.

## Authorship note

JBS, PK, and CTR are co-corresponding authors on this study.

