## Supplemental figure 1 for "Effect of obesity on the acute response to SARS-CoV-2 infection and development of post-acute sequelae of COVID-19 (PASC) in nonhuman primates"

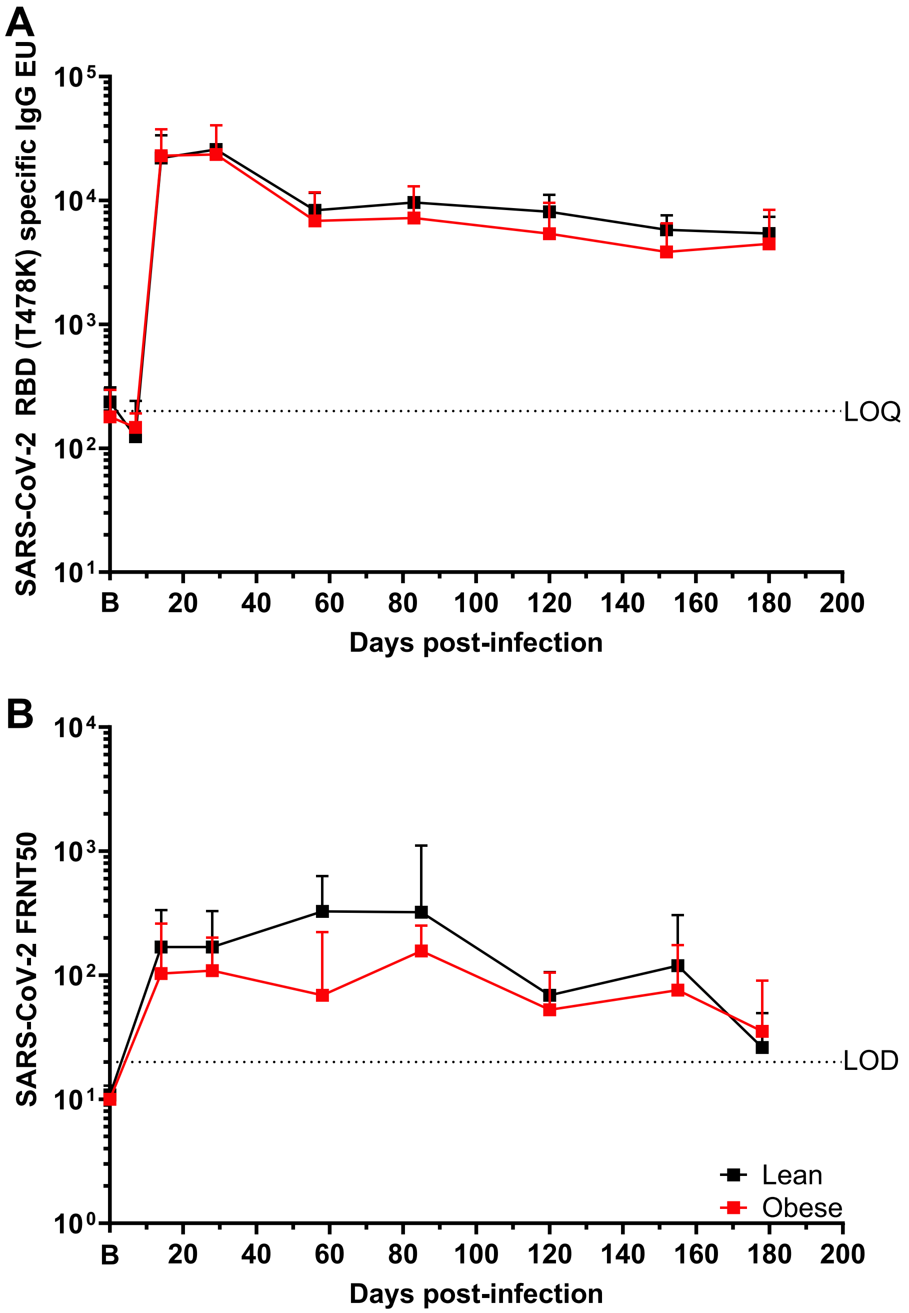


Supplemental figure 1. **SARS-CoV-2-specific serology in lean and obese animals.** A. RBD-binding IgG levels measured by ELISA. B. SARS-CoV-2-specific neutralizing Ab levels. All data are GMT +/-95% CI. LOQ: limit of quantitation. LOD: Limit of detection.
