## Supplemental figure 2 for "Effect of obesity on the acute response to SARS-CoV-2 infection and development of post-acute sequelae of COVID-19 (PASC) in nonhuman primates"

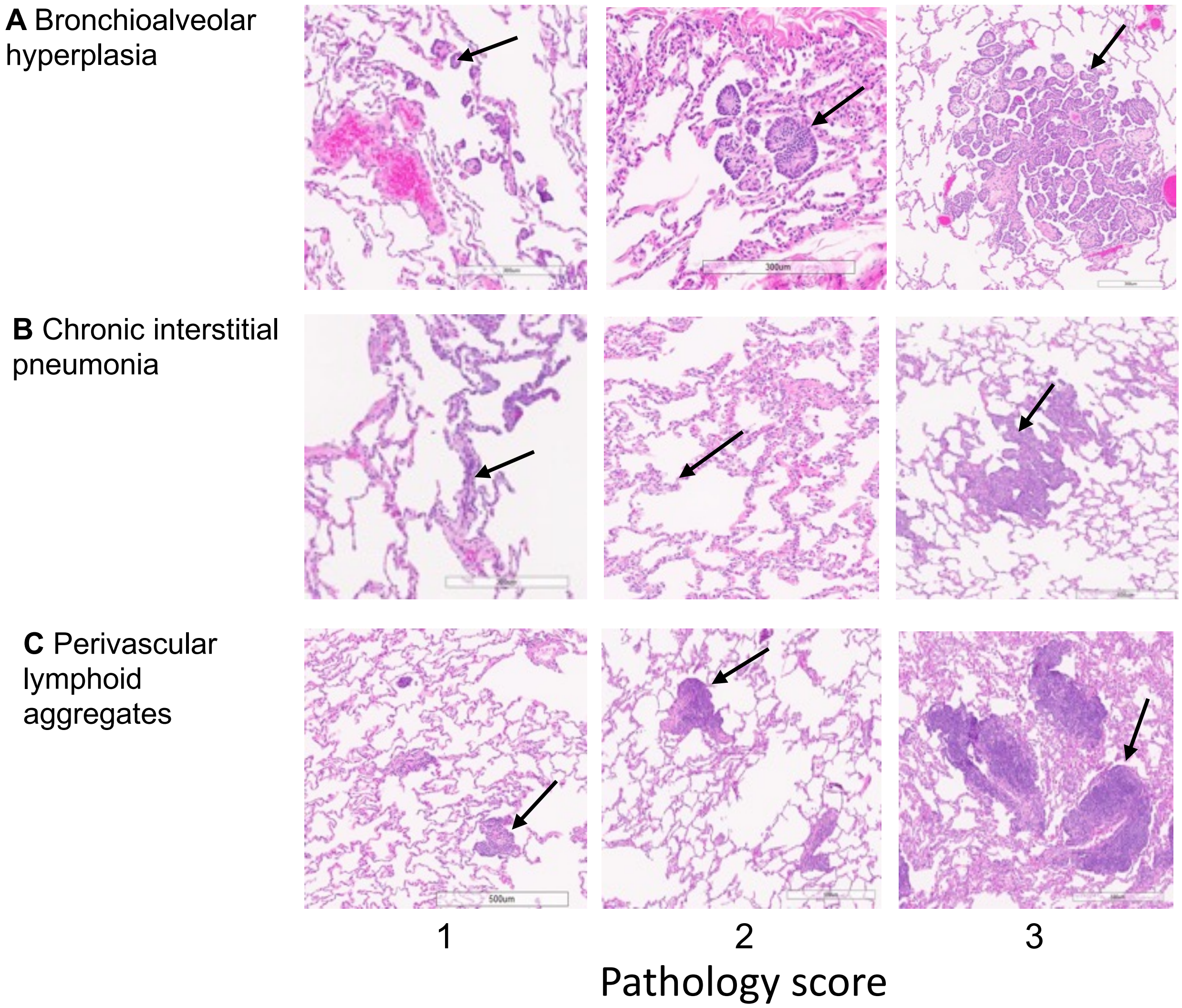


Supplemental figure 2. **Examples of microscopic lung lesions that reflect the histologic scoring criteria that are detailed in Table S1.** Scores range from 0 (no lesion, not pictured) up to a score of 3. A. Top row, bronchioloalveolar hyperplasia (arrows) is characterized by clusters of hyperplastic epithelial cells arising from the terminal bronchiole, alveolar duct and alveolus. B. Middle row, alveolar septa expanded with lymphocytes (arrows) are a feature of the chronic interstitial pneumonia observed here. C. Bottom row, variable numbers of lymphocytes surround multiple blood vessels (arrows). The extent of pathology featured here ranges from minimal to mild to moderate, corresponding to scores of 1, 2 or 3.
