## Supplemental figure 3 for "Effect of obesity on the acute response to SARS-CoV-2 infection and development of post-acute sequelae of COVID-19 (PASC) in nonhuman primates"

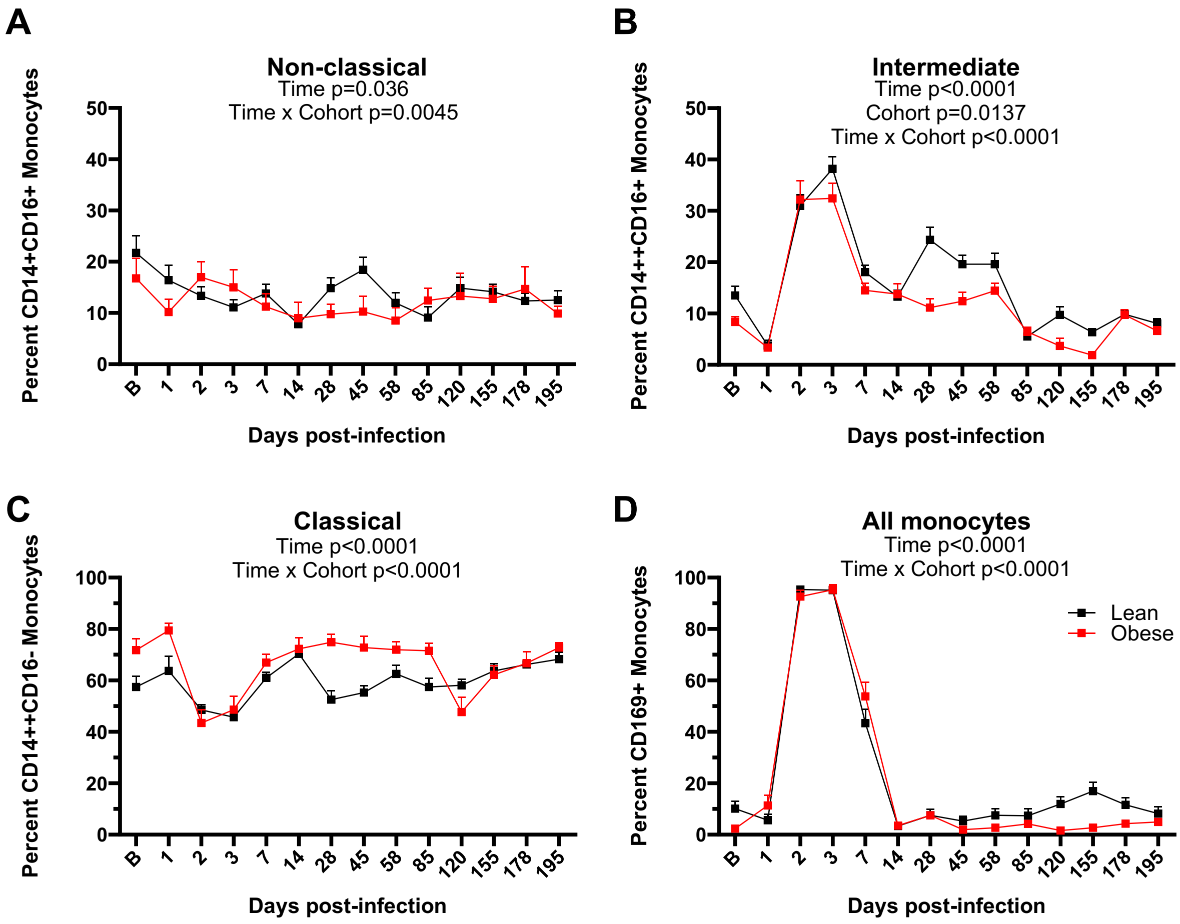


Supplemental figure 3. **Monocyte profiles in lean and obese animals.** PBMCs were analyzed by flow cytometry to distinguish non-classical (A), intermediate (B), and classical (C) subpopulations, as well as the CD169+ population (D) using the antibodies and gating strategies shown in Table S2 and Figure S7. All data are means ± SEM. Significance determined using mixed-effect analysis with Dunnett's post-hoc for multiple comparisons test.
