## Supplemental figure 4 for "Effect of obesity on the acute response to SARS-CoV-2 infection and development of post-acute sequelae of COVID-19 (PASC) in nonhuman primates"

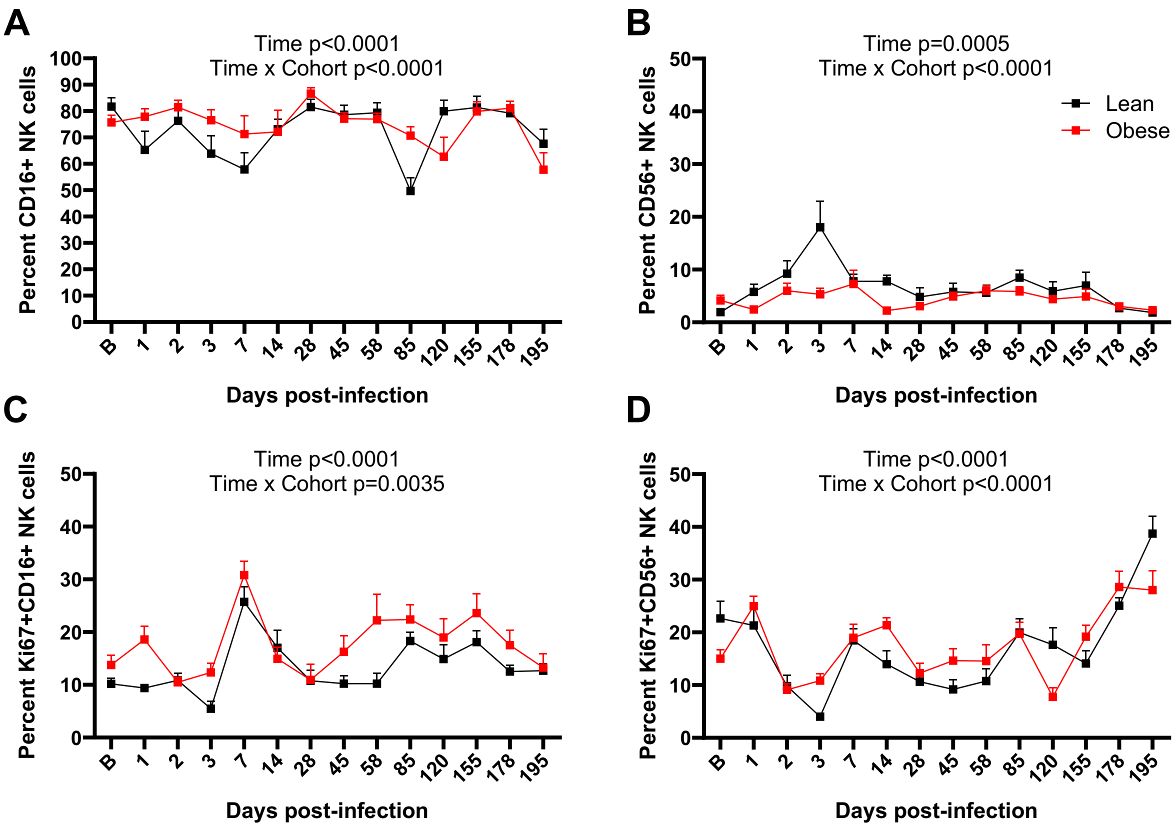


Supplemental figure 4. **NK cell profiles in lean and obese animals.** PBMCs were analyzed by flow cytometry to distinguish non-classical (A), intermediate (B), and classical (C) NK cell populations, as well as CD169+ NK cells (D) using antibodies and gating strategies shown in Supplemental Table 2 and Supplemental Figure 7. All data are means ± SEM. Significance determined using mixed-effect analysis with Dunnett's post-hoc for multiple comparisons test.
