## Supplemental figure 5 for "Effect of obesity on the acute response to SARS-CoV-2 infection and development of post-acute sequelae of COVID-19 (PASC) in nonhuman primates"

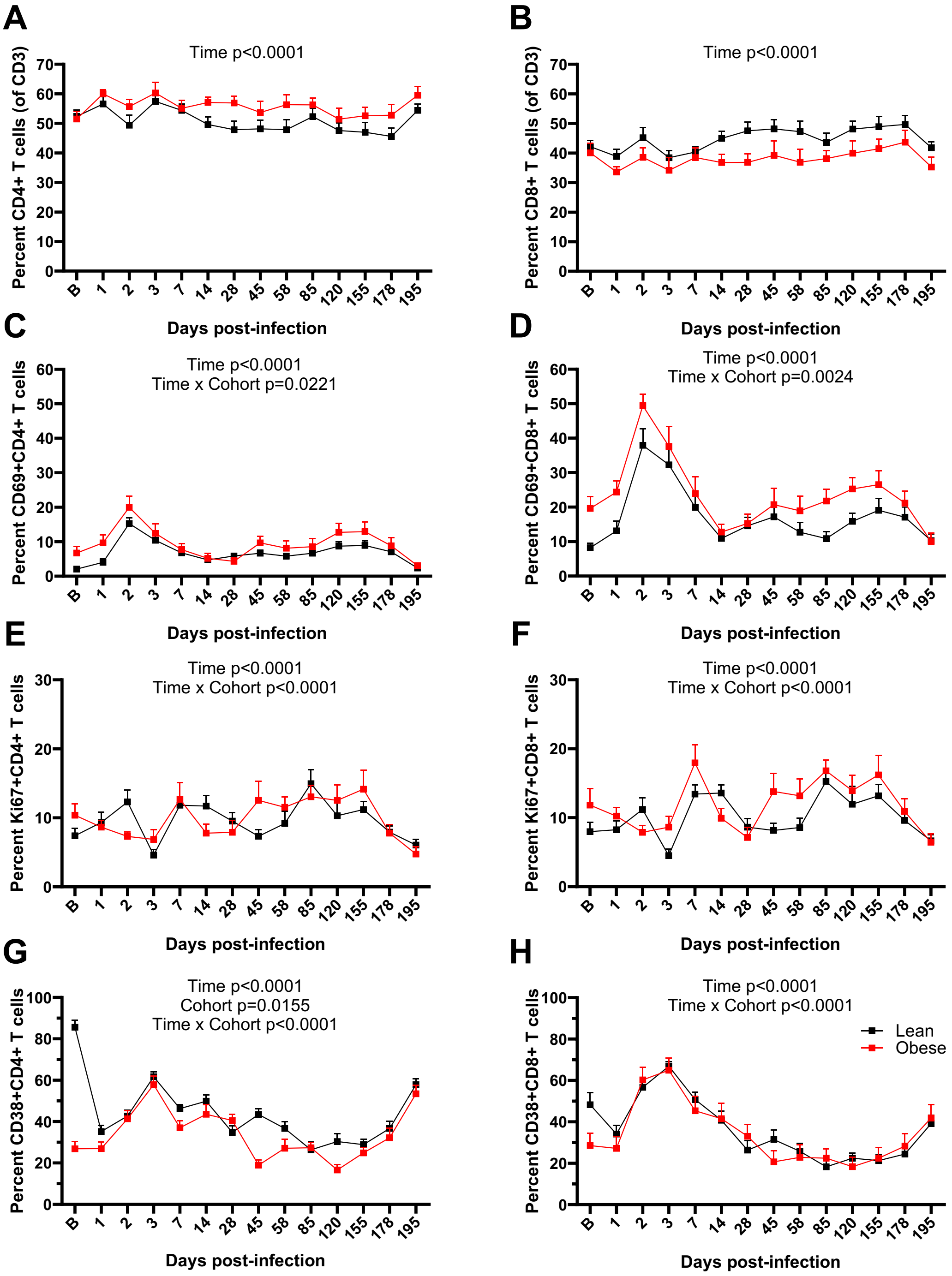


Supplemental figure 5. **T cell profiles in lean and obese animals.** PBMCs were analyzed by flow cytometry to distinguish CD4+ (A), CD8+ (B), CD69+CD4+ (C), CD69+CD8+ (D), Ki67+CD4+ (E), Ki67+CD8+ (F), CD38+CD4+ (G), and CD38+CD8+ (H) T cells using antibodies and gating strategies shown in Supplemental Table 2 and Supplemental Figure 7. All data are means ± SEM. Significance determined using mixed-effect analysis with Dunnett's post-hoc for multiple comparisons test.
