## Supplemental figure 6 for "Effect of obesity on the acute response to SARS-CoV-2 infection and development of post-acute sequelae of COVID-19 (PASC) in nonhuman primates"

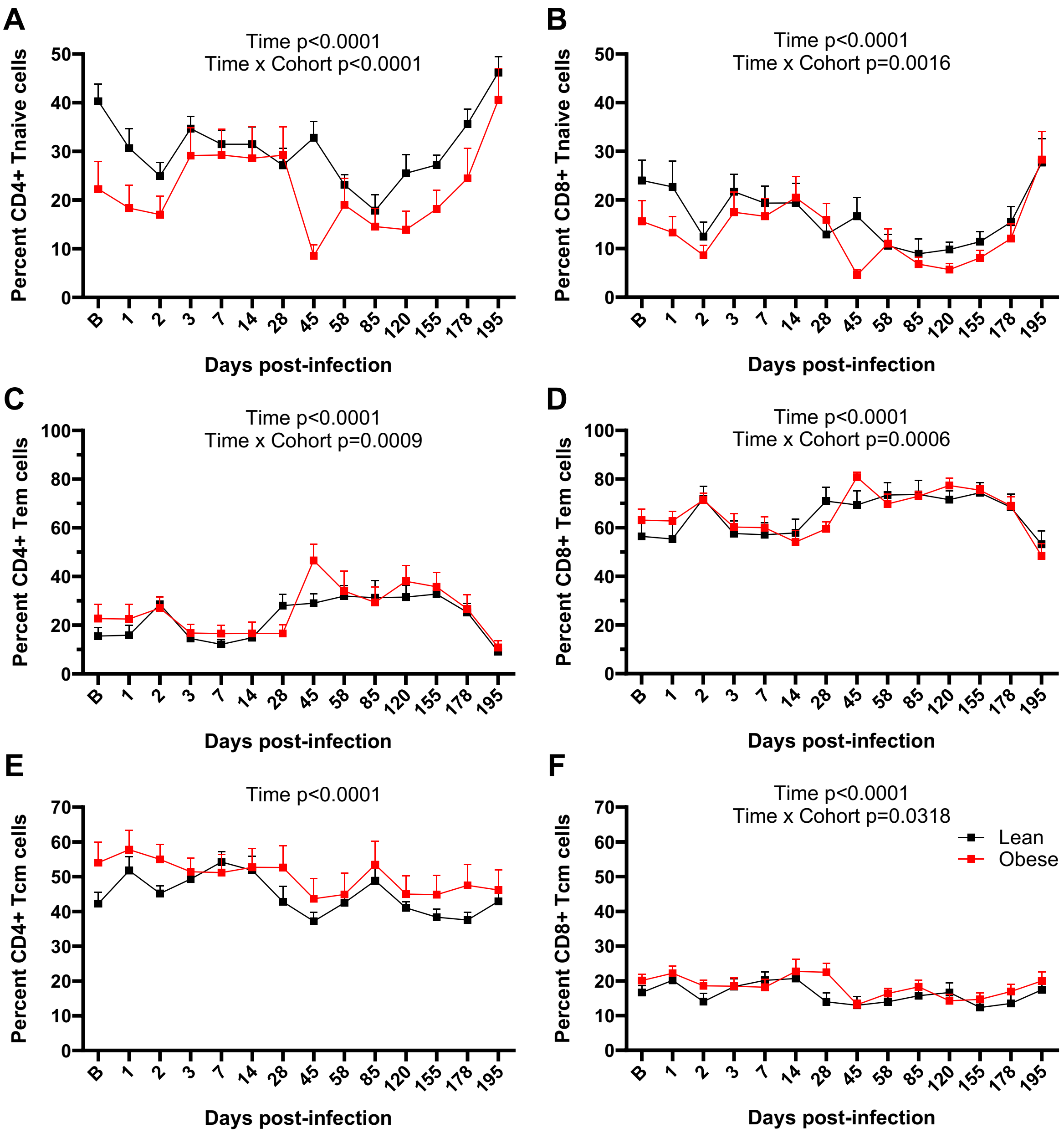


Supplemental figure 6. **T cell memory profiles in lean and obese animals.** PBMCs were analyzed by flow cytometry to distinguish CD4+ naive (A), CD8+ naive (B), CD4+ effector memory (Tem) (C), CD8+ Tem (D), CD4+ central memory (Tcm) (E), and CD8+ Tcm (F) T cells using antibodies and gating strategies shown in Supplemental Table 2 and Supplemental Figure 7. All data are means ± SEM. Significance determined using mixed-effect analysis with Dunnett's post-hoc for multiple comparisons test.
