## Supplemental figure 7 for "Effect of obesity on the acute response to SARS-CoV-2 infection and development of post-acute sequelae of COVID-19 (PASC) in nonhuman primates"

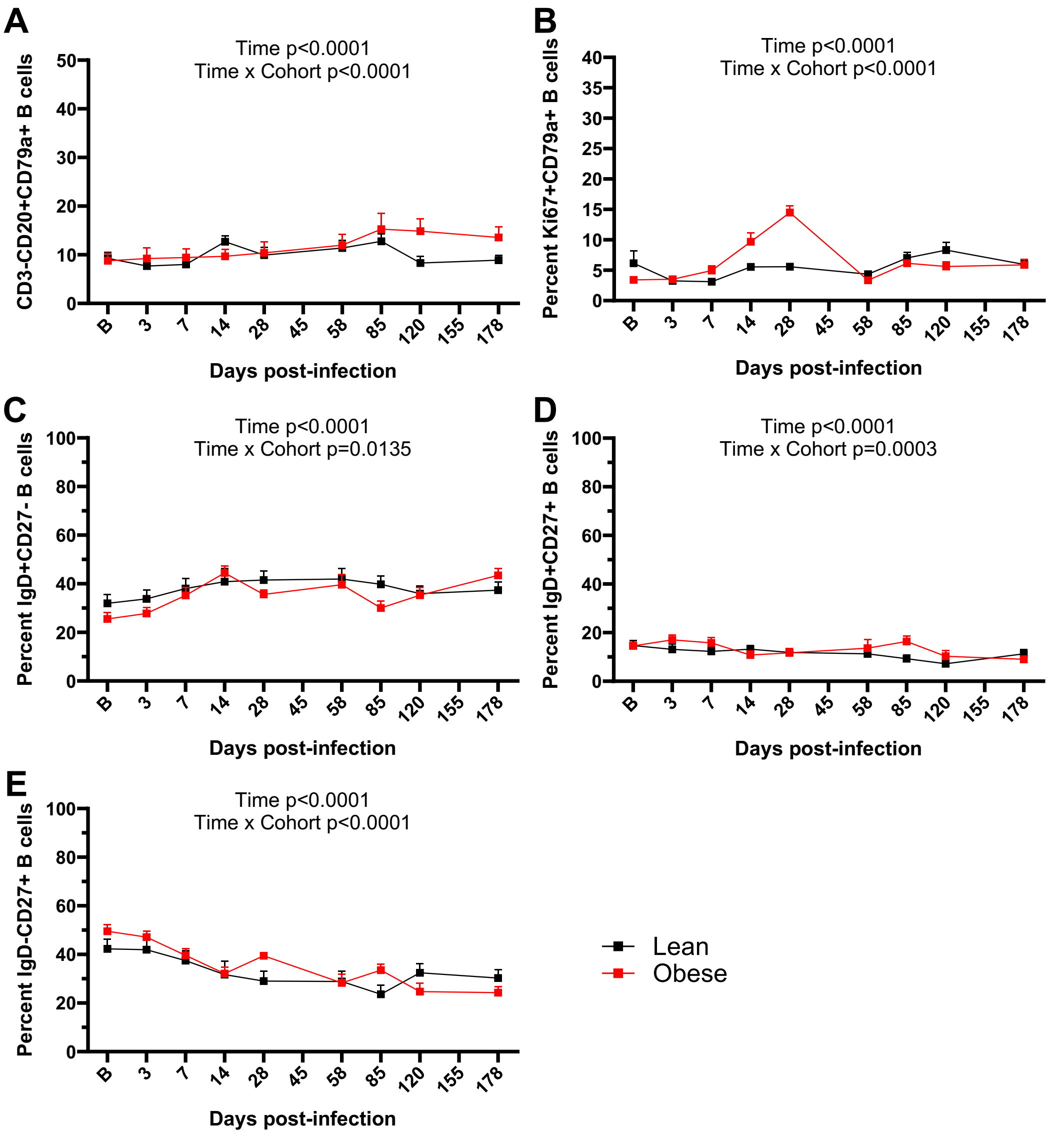


Supplemental figure 7. **B cell profiles in lean and obese animals.** PBMCs were analyzed by flow cytometry to distinguish total CD3-CD20+CD79a+ (A), activated Ki67+CD3-CD20+CD79a+ (B), naïve IgD+CD27- (C), IgM memory IgD+CD27+ (D), and class-switched memory IgD-CD27+ (E) B cells using antibodies and gating strategies shown in Supplemental Table 2 and Supplemental Figure 7. All data are means ± SEM. Significance determined using mixed-effect analysis with Dunnett's post-hoc for multiple comparisons test.
