## Supplemental figure 8 for "Effect of obesity on the acute response to SARS-CoV-2 infection and development of post-acute sequelae of COVID-19 (PASC) in nonhuman primates"

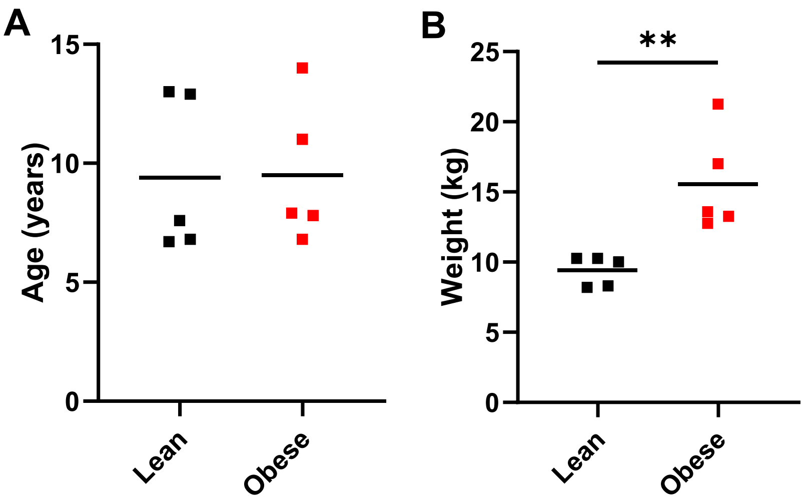


Supplemental figure 8. **Age (A) and weight (B) distribution in animals implanted with telemetry for BT and activity.** Significance was determined by unpaired 2-tailed t test. **, $p<0.01$.
