## Supplemental figure 9 for "Effect of obesity on the acute response to SARS-CoV-2 infection and development of post-acute sequelae of COVID-19 (PASC) in nonhuman primates"

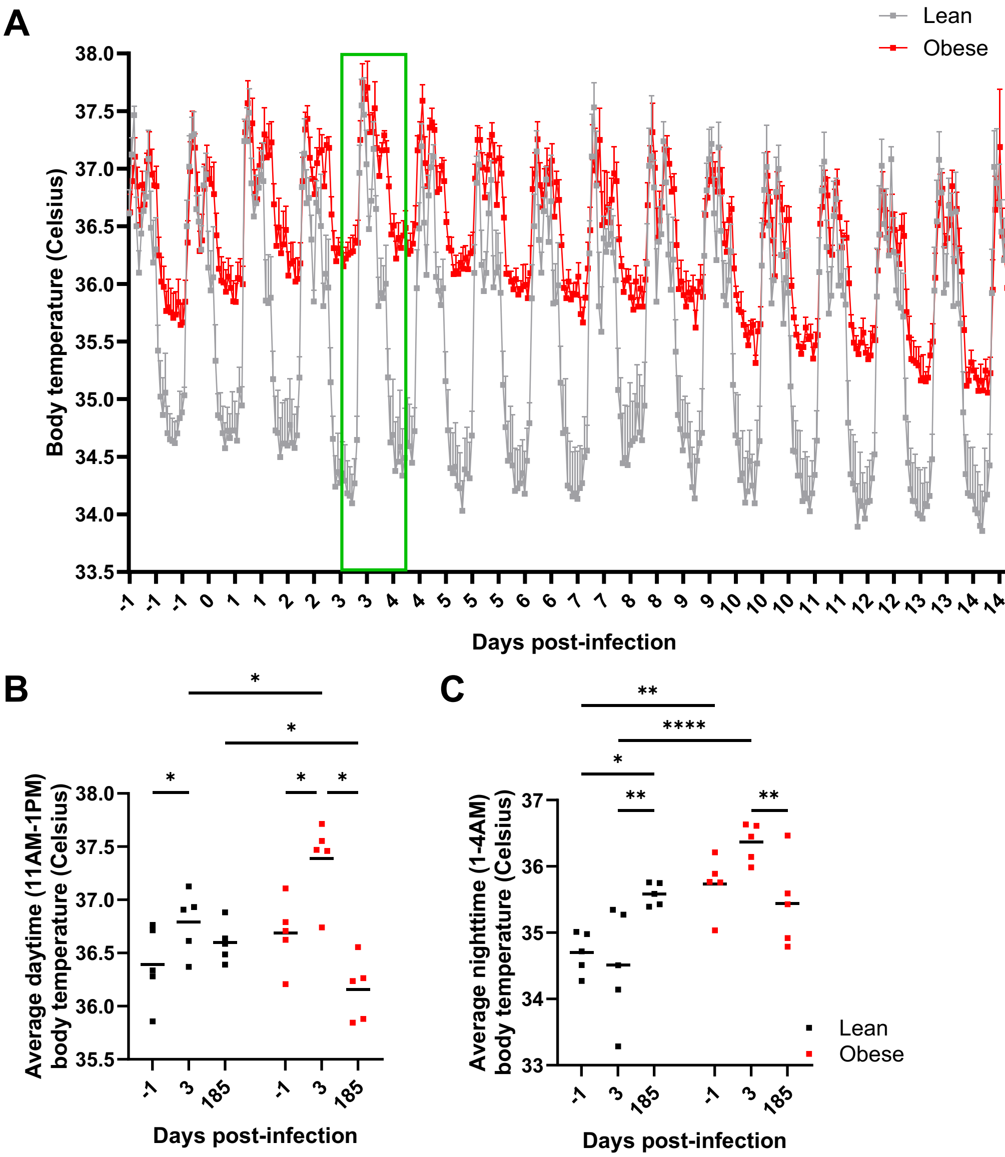


Supplemental figure 9. **Body temperature fluctuations following SARS-CoV-2 infection in lean and obese animals.** A. Circadian variation in BT in lean and obese animals from d -1 to 14 PI. Data are means ± SEM. B. Average daytime (11 AM-1 PM) and (C) nighttime (1-4 AM) BT at days -1, 3, and 185 PI. Significance determined by 2-way ANOVA w/Tukey's multiple comparison test. *, $p<0.05$; **, $p<0.01$; ****, $p<0.0001$.
