## Supplemental figure 10 for "Effect of obesity on the acute response to SARS-CoV-2 infection and development of post-acute sequelae of COVID-19 (PASC) in nonhuman primates"

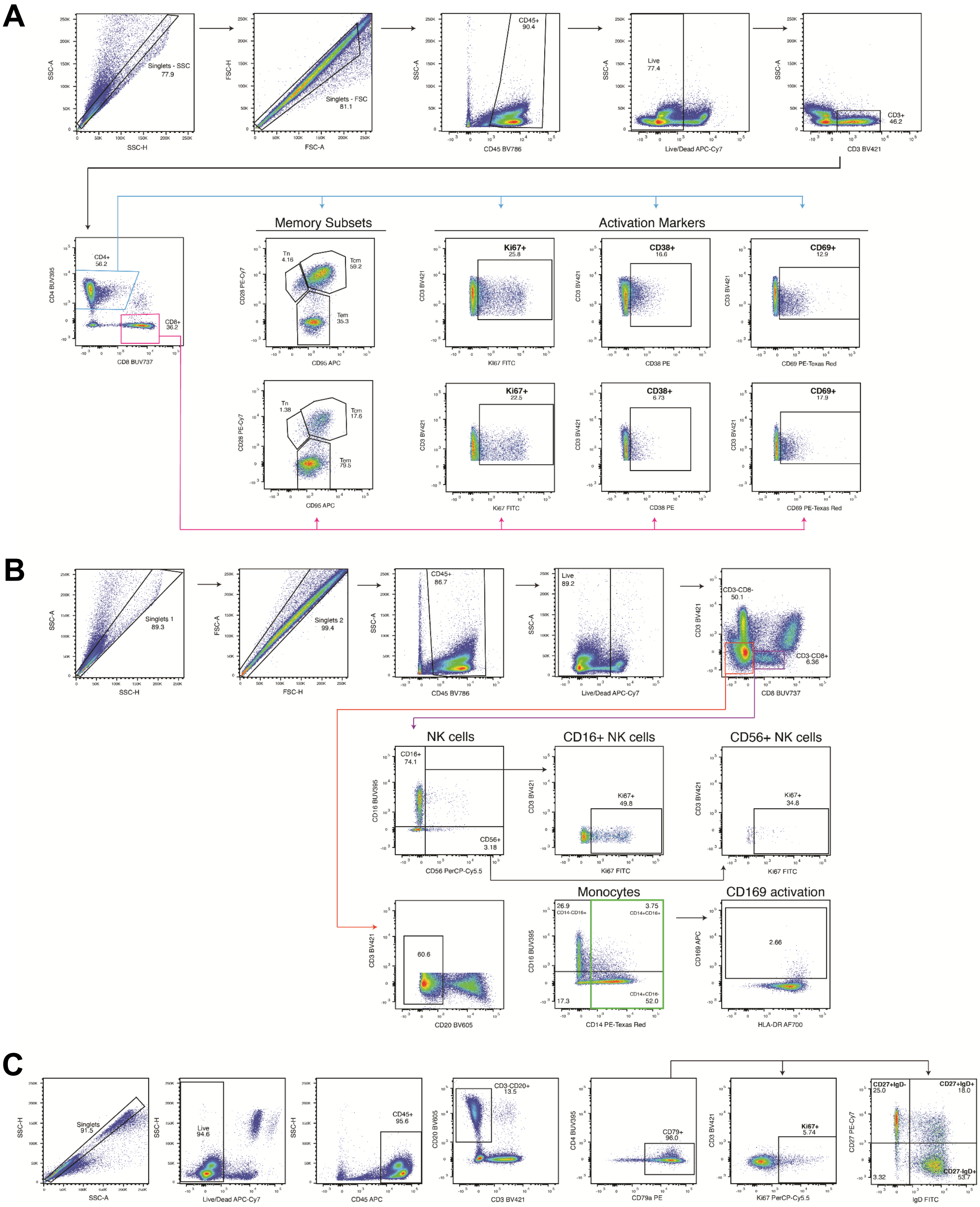


Supplemental figure 10. **Gating strategies for flow cytometry analyses.** PBMCs were characterized for T cells (A), monocytes and NK cells (B), and B cells (C).
