## Supplemental table 1 for "Effect of obesity on the acute response to SARS-CoV-2 infection and development of post-acute sequelae of COVID-19 (PASC) in nonhuman primates"

| Score | 0 | 1 | 2 | 3 |
| --- | --- | --- | --- | --- |
| **Bronchioalveolar hyperplasia** | None | 1 to 5 foci,<br>all <500 μM | 6 to 10 foci,<br>all <500 μM | >10 foci and/or at least one focus >500 μM |
| **Lymphocytic interstitial pneumonia** | None | 1 to 5 foci | 6 or more foci | At least one focus > 1mm |
| **Perivascular lymphoid aggregates** | None | Minimal | Mild | Moderate |

Supplemental table 1. **Lesions used to determine lung pathology score.** Nine lung slides, including two slides of the caudal lobes and one of the other lung lobes, were scanned at 40x with an Aperio AT2 microscope slide scanner (Leica Biosystems, Deer Park, IL). A scoring system was developed, and histologic findings were then recorded and aggregated. Those findings considered to represent responses to chronic injury that occurred in multiple animals were used as the basis of subsequent scoring illustrated in the table above. Specifically, bronchioloalveolar hyperplasia, lymphocytic interstitial pneumonia, and perivascular lymphoid aggregates were selected as being potentially associated with SARS-CoV-2 infection. A score ranging from 0 to 3 was assigned for each of these three histologic features identified in every lung lobe examined of individual animals and were added together for a final score.
