## Supplemental table 2 for "Effect of obesity on the acute response to SARS-CoV-2 infection and development of post-acute sequelae of COVID-19 (PASC) in nonhuman primates"

| Antibody Target | Fluorophore | Clone | Host Species | Source | Catalog # |
|---|---|---|---|---|---|
| CD4 | BUV395 | L200 | Mouse | BD Biosciences | 564107 |
| CD8 | BUV737 | SK1 | Mouse | BD Biosciences | 612754 |
| CD45 | BV786 | D058-1283 | Mouse | BD Biosciences | 563861 |
| CD28 | PE-Cy7 | CD28.2 | Mouse | BD Biosciences | 560684 |
| CD3 | PB | SP34-2 | Mouse | BD Biosciences | 558124 |
| CD95 | APC | DX2 | Mouse | BioLegend | 305612 |
| CD38 | PE | OKT10 | Mouse | NHPRR | PR-3802 |
| HLA-DR | AF700 | G46-6 | Mouse | BD Biosciences | 560743 |
| CD69 | PE-TR | FN50 | Mouse | BioLegend | 310942 |
| Ki67 | FITC | B56 | Mouse | BD Biosciences | 556026 |
| CD20 | BV605 | 2H7 | Mouse | BD Biosciences | 747736 |
| CD14 | ECD | RMO52 | Mouse | Beckman Coulter | IM2707U |
| CD56 | PerCP-Cy5.5 | B159 | Mouse | BD Biosciences | 560842 |
| CD16 | BUV395 | 3GB | Mouse | BD Biosciences | 563785 |
| CD169 | APC | 7-239 | Mouse | Biolegend | 346008 |
| IgD | FITC | - | Goat | Southern Biotech | 2030-09 |
| CD45 | APC | D058-1283 | Mouse | BD Biosciences | 561290 |
| CD27 | PE-Cy7 | O323 | Mouse | eBioscience | 25-0279-42 |
| CD79a | PE | HM47 | Mouse | eBioscience | 12-0792-42 |
| Ki67 | PerCP-Cy5.5 | B56 | Mouse | BD Biosciences | 561284 |
| Live/Dead ARD | APC-Cy7 | - | - | Life Technologies | 34959 |

Supplemental table 2. **Antibodies used in flow cytometry.**
